# Living on the edge: warmer climate reduces leaf thermal safety margins and causes gas exchange decoupling in Mediterranean shrubs

**DOI:** 10.64898/2025.12.06.692718

**Authors:** Margaux Didion-Gency, Pol Soler-Ruiz, Eva Castells, Jordi Martinez-Vilalta

## Abstract

Global warming pushes plants close to their leaf thermal limits, altering carbon uptake, growth and survival. Therefore, investigating leaf thermal tolerance adjustment is essential to understand future vegetation dynamics, especially in high-risk Mediterranean shrubs facing hot and dry summers.

We measured dark-adapted leaf fluorescence (F_v_/F_m_), thermal thresholds (T_crit_, T_50_, T_max_), optimal assimilation temperature (T_opt_), thermal safety margin (TSM), leaf and air- temperatures (T_air_, T_leaf_) and gas exchange (A, g_s_, E) in six shrub species in six sites along a climatic gradient in Catalonia, Spain.

We found that T_opt_ increased as conditions warmed, while F_v_/F_m_ and T_max_ showed quadratic responses decreasing in the warmest sites. Hence, TSM declined and approached 0 at the hottest sites, indicating that shrubs were operating close to their thermal limits. Warmer T_air_ under high levels of solar radiation raised T_leaf_, though some species maintained T_leaf_ < T_air_ during daytime, suggesting a passive or active leaf cooling at high temperatures exceeding solar energy. Moreover, E remained high despite low A at higher T_leaf_, revealing a decoupling of gas exchange at high temperatures.

Incorporating leaf thermal tolerance adjustments and gas exchange decoupling at extreme temperatures into vegetation models could improve predictions of shrub function and dynamics under warmer climates.

**Highlight:** Our study highlights that Mediterranean shrubs have limited capacity to adjust leaf thermal tolerance at warmer sites, resulting in narrower thermal safety margins.

The decrease in leaf temperatures through evaporative cooling may not compensate for the more frequent and intense heatwaves, leading to irreversible damage.

These findings highlight the importance of incorporating leaf thermal tolerance adjustment and gas exchange decoupling into future vegetation models.

## Introduction

Global rising air temperatures and more frequent heatwaves are expected under climate change, with a business-as-usual scenario predicting a temperature increase of 4.4 °C by 2100 (IPCC, 2023). The rise in air temperature is expected to reduce plants carbon assimilation (Ciais *et al*., 2005; Rödenbeck *et al*., 2020; Didion-Gency *et al*., 2022), increase widespread leaf damage (Teskey *et al*., 2015), and induce shifts in species distributions, particularly at the species’ rear edge (Allen *et al*., 2015; Brodribb *et al*., 2020). Leaf thermal tolerance traits can be used to identify temperatures that induce plant irreversible physiological damage (Aspinwall *et al*., 2019; Lancaster & Humphreys, 2020). Their use has expanded significantly in recent decades, enabling comparisons of thermal tolerance both between and within species and to predict leaf thermal safety margins under warmer climates (Deutsch *et al*., 2008; Zhu *et al*., 2018; Rao *et al*., 2023; Gauthey *et al*., 2023).

While most studies on leaf-level physiological thermal tolerance have focused on tropical trees, other woody species, such as shrubs, remain comparatively understudied. Yet, shrubs dominate many arid and semi-arid ecosystems and are projected to expand as conditions get warmer and drier (Batllori *et al*., 2020). Moreover, the Mediterranean region have been identified as a prominent climate response hotspots (Giorgi, 2006), with projections under business-as-usual scenario indicating summer temperature increases of up to 6 °C and precipitation decreases of up to 35% by 2100 (Giorgi *et al*., 2004). Consequently, Mediterranean shrub species, in particular, are projected to replace numerous existing forests, and given their exposure to hot and dry summers, may experience some of the most severe climate change impacts (Peñuelas *et al*., 2017; Peñuelas & Sardans, 2021).

Leaf thermal tolerance can be assessed by measuring functional thermal thresholds of photosynthetic efficiency of photosystem II (PSII; F_v_/F_m_; Cook *et al*., 2024; Didion-Gency *et al*., 2025), regarded as the most heat-sensitive component of the photosynthetic machinery (Berry & Bjorkman, 1980; Maxwell & Johnson, 2000a). High-temperature leaf thermal thresholds include the critical temperature that induces an abrupt decrease of F_v_/F_m_ (T_crit_; Slot *et al*., 2021), the temperature causing a 50 % reduction of F_v_/F_m_ (T_50_; Curtis *et al*., 2014; Ahrens *et al*., 2021), and maximum temperature under which F_v_/F_m_ approaches 0 (T_max_; Didion-Gency *et al*., 2025). Recent studies suggests that species can adjust their thermal thresholds in responses to warmer climates in order to increase their long-term survival (Slot *et al*., 2021). Thus, species from hotter climates generally exhibit higher thermal thresholds than those from cooler ones (Zhu *et al*., 2018). For instance, O’sullivan *et al*. (2017) reported T_crit_ ranging from 43.5 to 50.8 °C, and T_max_ from 51.0 to 60.6 °C for polar and tropical species, respectively. Similarly, Knight & Ackerly (2002) observed T_50_ ranging from 46.4 to 55.3 °C, and T_max_ from 50.2 to 64.1 °C for Mediterranean and desertic species, respectively. Furthermore, individual species may also adjust to heat by increasing their thermal threshold when growing in warmer sites (Feeley *et al*., 2020). For instance, Slot *et al*., (2021) observed a +0.6 °C and +0.4 °C increase in T_crit_ and T_50_, respectively, per 1 °C increase mean annual temperature on tropical species. Furthermore, studies showed that exposure to higher growth temperatures can induce an accumulation of heat-shock proteins, which can temporarily mitigate possible physiological thermal damages (Wahid *et al*., 2007; Eaton-Rye *et al*., 2012; Aspinwall *et al*., 2019), thereby raising short-term thermal tolerance. However, the extent to which leaf thermal thresholds can adjust in species grown at warmer climate, such as Mediterranean shrubs, and whether such adjustment can fully compensate for the projected increase in temperature in Mediterranean climates, remains an important uncertainty, particularly under natural conditions.

In addition to PSII thermal thresholds, thermal acclimation can be determined by measuring the adjustment of the leaf optimal temperature (T_opt_) that allows optimal assimilation (A_opt_) under warm temperatures. Recent studies suggested that species can also adjust their T_opt_ and A_opt_ thresholds in responses to warmer climates, where species from hotter climates generally exhibit a higher T_opt_ than those from cooler ones (Tan *et al*., 2017; Chen *et al*., 2021). For instance, Crous *et al*. (2022) observed a +0.34 °C increase in T_opt_ per 1 °C increase in mean temperature of the warmest quarter on boreal to tropical species. Similarly, Sendall *et al*. (2015) observed an increase in T_opt_ in years with higher temperatures for both boreal and temperate species. Moreover, it has been shown that T_opt_ generally ranges from 20 to 30 °C, but assimilation (A) can remain stable up to 46 °C in high thermal tolerant species (Downton *et al*., 1984). Irreversible damage to the photosynthetic machinery is typically observed between 45 to 50 °C (Krause *et al*., 2010; Hüve *et al*., 2019), corresponding approximately to T_50_. The decline in A above T_opt_ may result from the coupled limitations of gas exchange, including a decrease in stomatal conductance (g_s_) due to stomatal closure in response to increased vapor pressure deficit (Kumarathunge *et al*., 2020; Grossiord *et al*., 2020), as well as alteration of the photosynthetic biochemistry, including higher degradation of the Rubisco activase (Bernacchi *et al*., 2002; Sage & Kubien, 2007), lower regeneration of ribulose bisphosphate (Kubien & Sage, 2008), and lower electron transport rate (Wise *et al*., 2004), or from higher leaf respiration rates (Lin *et al*., 2012).

Recent works have also demonstrated that g_s_ and transpiration (E) have a higher thermal tolerance than A by about 10 °C (Didion-Gency *et al*., 2025), leading to a decoupling of A and g_s_ or E at high temperatures (T > T_opt_; Diao *et al*., 2024; Gauthey *et al*., 2024). During this phase of photosynthetic machinery impairment (T_opt_ < T < T_crit_), it has been hypothesized that the maintenance of low but non-zero g_s_ and continued E at higher temperatures could be associated with an active and/or passive evaporative cooling, resulting in a lower leaf temperature (T_leaf_) to help dissipate excess heat, prevent thermal damage, and allow leaf survival for future carbon gain (Schymanski *et al*., 2013; Marchin *et al*., 2016; Urban *et al*., 2017; Drake *et al*., 2018a). Thus, another potential adjustment to improve long-term survival under warmer climate would be to reduce the maximum T_leaf_ experienced (T_leaf, max_) through evaporative cooling. However, whether the residual E observed at high temperatures results from an active incomplete stomatal closure or to passive stomatal and/or cuticular leakage remains unclear.

At high temperatures, leaf water loss may continue even after stomatal closure, through residual stomatal leakage and cuticular conductance, known as leaf residual conductance (g_res_), or in some cases attributed solely to cuticular conductance (g_min_; Duursma *et al*., 2019; Fernandes *et al*., 2025). Studies observed that individuals experiencing higher temperatures generally exhibit higher g_min_ than those from cooler ones (Garen & Michaletz, 2025). For instance, Schuster *et al*. (2016) observed a x2.4 factor increase in *g*_min_ from 15 to 50 °C on desertic species. Similarly, Bueno *et al*. (2019) observed a x3.2 factor increase in g_min_ from 25 to 50 °C, also on desertic species. Recent studies suggested that this water loss after stomatal closure at high temperatures can contribute to plant hydraulic damage (Schönbeck *et al*., 2022). Nevertheless, a reduction of T_leaf_ through evaporative cooling may lead to higher thermal safety margin (TSM), defined as the difference between T_crit_ and T_leaf,max_ (Leon-Garcia & Lasso, 2019). Ideally, this mechanism would allow plants to maintain TSM > 0. However, only a few studies have examined at TSM along natural climatic gradients, but these showed that species growing in hotter climates may still generally exhibit lower TSM than those from cooler ones (Kullberg *et al*., 2024), because thermal adjustments are not able to fully compensate for higher exposure, resulting in more vulnerable individuals.

In this study, we aimed to understand how six Mediterranean shrub species (i.e., *Amelanchier ovalis* Medik., *Arbutus unedo* L., *Buxus sempervirens* L., *Pistacia lentiscus* L., *Rhamnus alaternus* L., and *Salvia rosmarinus* Spenn.) adjusted their leaf thermal tolerance, gas exchange, stomatal opening, and thermal safety margin at six sites along a climatic gradient. Thus, our objectives were to (1) assess the adjustment of leaf thermal tolerance traits (i.e., F_v_/F_m_, T_crit_, T_50_, T_max_, T_opt_, A_opt_, TSM) as climate warms, and to (2) investigate whether these species exhibit a pattern of gas exchange decoupling (i.e., A *vs*. gs and E) at high temperatures. We expected that (1) shrub species growing under warmer climates will have traits associated with higher thermal tolerance compared to the ones growing in cooler ones, but not necessarily sufficient to compensate for the higher heat exposure, leading to lower TSM; (2a) plants of all species will exhibit gas exchange decoupling at high temperatures due to continued transpiration (while photosynthesis declines to near zero) as a mechanism to reduce T_leaf_ through evaporative cooling and avoid critical overheating, and (2b) shrub species growing in warmer climates will exhibit a more pronounced decoupling.

## Materials and methods

### Study sites and sampling design

The study was conducted at six sites along a climatic gradient in Catalonia, NE Spain (Fig. **1**; Table **1**). The warmest and driest site was Pas de l’Ase (PAS), followed by Alòs de Balaguer (ALO), Garraf (GAR), Cingles de Bertí (CIN), Montsec (MON), and Pentina (PEN) as the coolest and wettest site (Fig. **1**, Table **1**). All sites correspond to open Mediterranean shrublands with full sun exposure, located in natural forests area. Six shrub species (*Amelanchier ovalis* Medik., *Arbutus unedo* L., *Buxus sempervirens* L., *Pistacia lentiscus* L., *Rhamnus alaternus* L., and *Salvia rosmarinus* Spenn.) were selected. The environmental gradient studied encompasses either their full climatic range (*A. ovalis*, *B. sempervirens*), core range (*A. unedo*, *P. lentiscus*), or central to dry range (*R. alaternus*, *S. rosmarinus*; Fig. **S1**). These species are among the most common plants in each site. Other abundant woody species in the study sites were *Quercus coccifera* L., *Buplerum fructicosum* L., *Erica multiflora* L., *Ulex parviflorus* Pourr., and *Thymus vulgaris* L. To limit confounding effects, only sites without signs of recent management or major disturbances were selected. Each species was represented in at least three of the six sites (Table **1**), covering a substantial part of their climatic distribution (Fig. **S1**). In each site, seven to nine shrubs per species were randomly selected, with a total of 174 individuals, and characterized by their height (H), basal diameter (BD), stem number (N_stem_), plant competition (i.e., plant cover within a circular band with the width equal to the height of the target individual, P_competition_,), and individual plant crown area (P_crown_; Table **S1**). Plant crown area (P_crown_) for each individual was calculated as follows:

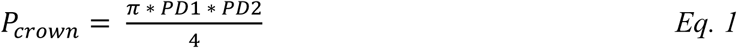

where PD1 represents the largest plant diameter, and PD2 the perpendicular diameter to PD1.

**Figure 1:**
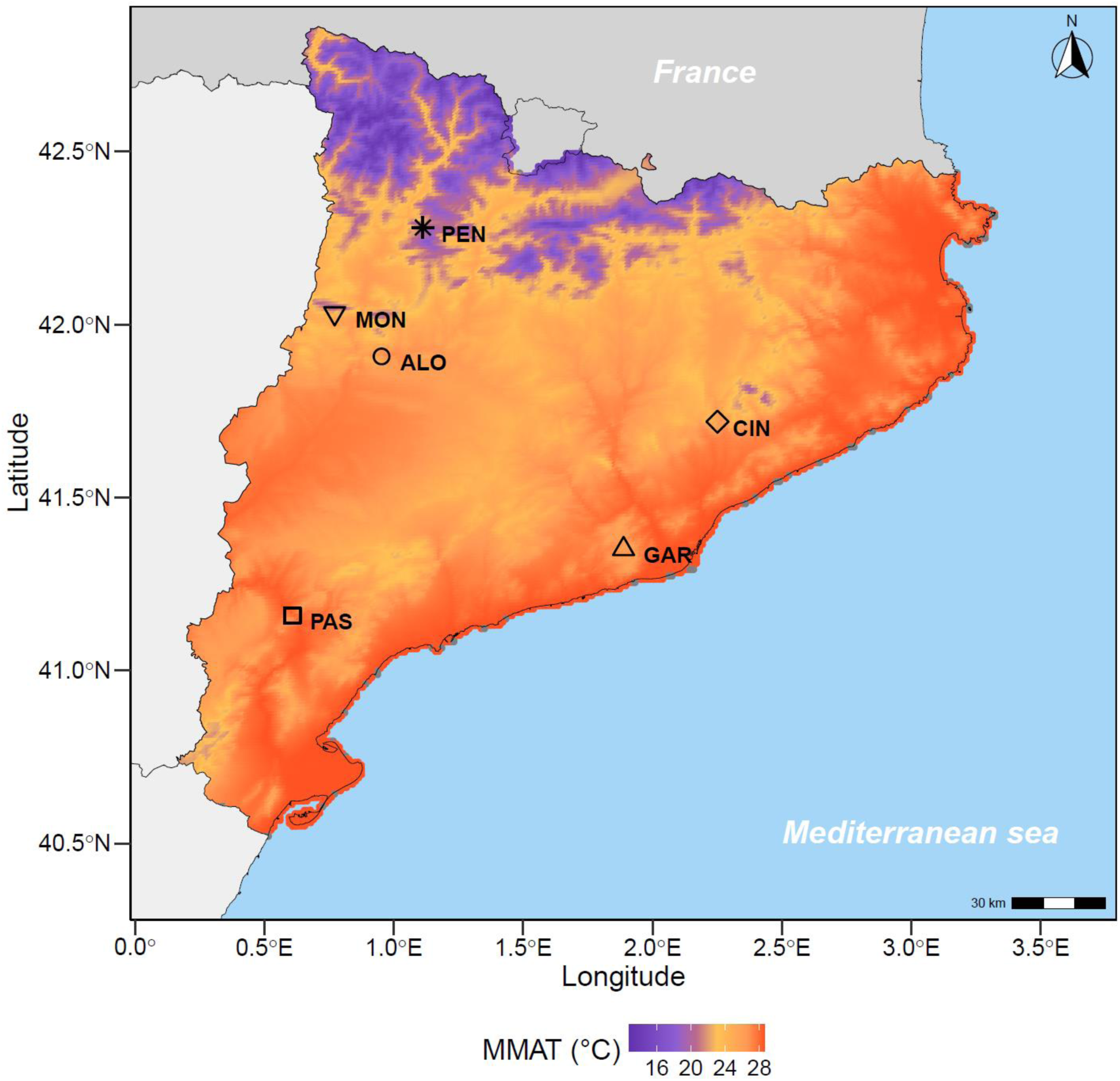
Geographical locations of the study sites (i.e., PAS: Pas de l’Ase, ALO: Alòs de Balaguer, GAR: Garraf, CIN: Cingles de Bertí, MON: Montsec, PEN: Pentina) in Catalonia (NE, Spain) located in a map of the maximum monthly air temperature (MMAT) for 2024. The color gradients correspond to the temperature gradient where dark purple and dark orange represent the cooler and warmer areas, respectively.

**Table 1:** Characteristics of the study sites.

| Location | Site<br>code | Species | Mean<br>height<br>(cm) | Total<br>cover<br>(%) | Lat<br>(°N) | Lon<br>(°E) | Total<br>area<br>(ha) | Elevation<br>(m a.s.l) | MMAT<br>2009 - 2024<br>(°C) | MMAT<br>2024<br>(°C) | T<sub>air, min</sub><br>2024<br>(°C) | T<sub>air, max</sub><br>2024<br>(°C) | Summer<br>P/PET<br>1951 - 1999 |
| --- | --- | --- | --- | --- | --- | --- | --- | --- | --- | --- | --- | --- | --- |
| Pas de l'Ase | PAS | Au, Pl, Ra, Sr | 183.94 | 78.26 | 41.16 | 0.61 | 0.79 | 106 | 27.82 | 28.04 | -0.85 | 38.93 | 0.22 |
| Alòs de Balaguer | ALO | Ao, Bs, Ra, Sr | 150.50 | 85.31 | 41.91 | 0.95 | 0.36 | 431 | 26.04 | 26.26 | -2.23 | 36.93 | 0.31 |
| Garraf | GAR | Au, Bs, Pl, Ra, Sr | 154.77 | 84.85 | 41.35 | 1.89 | 0.29 | 584 | 24.31 | 25.41 | -2.88 | 35.98 | 0.47 |
| Cingles de Bertí | CIN | Au, Pl, Sr | 171.92 | 84.64 | 41.72 | 2.25 | 0.17 | 639 | 23.91 | 25.12 | -3.11 | 35.65 | 0.55 |
| Montsec | MON | Ao, Bs, Ra, Sr | 163.38 | 74.52 | 42.03 | 0.77 | 3.24 | 1298 | 20.99 | 21.49 | -6.59 | 31.67 | 0.66 |
| Pentina | PEN | Ao, Bs | 281.65 | 57.72 | 42.28 | 1.11 | 0.13 | 1526 | 18.78 | 20.25 | -7.91 | 30.48 | 0.91 |

<sup>1</sup>Abbreviation: Ao: *A. ovalis*, Au: *A. unedo* , Bs: *B. sempervirens*, Pl: *P. lentiscus*, Ra: *R. alaternus*, and Sr: *S. rosmarinus*, Total cover: total plant canopy cover, Lat: latitude, Lon: longitude, MMAT: maximum monthly air temperature, T: temperature, P/PET: maximum monthly precipitation divided by potential evapotranspiration.
<sup>2</sup>Temperatures values were obtained from Meteo.cat stations where site-specific downscaling was performed.
<sup>3</sup>P/PET values were obtained from Catalan Atlas and site-specific calculated as the total precipitation from April to September period divided by the total potential evapotranspiration over the same period. Lower values indicate higher aridity.

Fourteen physiological traits related to leaf thermal tolerance were obtained once on all plants between mid-July and mid-August 2024, corresponding to the summer period in Mediterranean region, characterized by high temperatures and low soil moisture levels (Table **2**). Measurements started from the warmer and drier sites and finished in the cooler and wetter ones to account for phenological differences among areas, except for PAS where the gas exchange temperature response curves were repeated at the end of the measurement campaign due to technical issues during the initial campaign (see below). Measurements and samples were obtained under sunny conditions (i.e., daily mean PPDF ≈ 1150 μmol m⁻² s⁻¹), with experienced environmental conditions generally consistent with those of the gradient sites (Table **S2**).

**Table 2:** List of the leaf thermal tolerance traits measured.

| Traits | Symbol | Units | Measurement type | Description |
| --- | --- | --- | --- | --- |
| Photosynthetic efficiency of PSII | $F_v/F_m$ | - | Single point on *in-situ* shoots | Efficiency of light energy conversion for assimilation |
| Critical leaf temperature that induces an abrupt increase in $F_v/F_m$ | $T_{crit}$ | °C | Curves derived on cut shoots | Temperature threshold at which irreversible damage to PSII begins |
| Temperature causing a 50% reduction of $F_v/F_m$ | $T_{50}$ | °C | Curves derived on cut shoots | Temperature threshold at which PSII efficiency drops by 50% |
| Maximum temperature under which $F_v/F_m$ approaches 0 | $T_{max}$ | °C | Curves derived on cut shoots | Temperature threshold at which PSII efficiency drops to ~ 0 |
| Leaf assimilation | A | $\mu\text{mol m}^{-2} \text{s}^{-1}$ | Curves derived on cut shoots | Rate of carbon uptake |
| Leaf stomatal conductance | $g_s$ | $\text{mmol m}^{-2} \text{s}^{-1}$ | Curves derived on cut shoots | Measure of stomatal opening, controlling gas exchange and water loss |
| Leaf transpiration | E | $\text{mol m}^{-2} \text{s}^{-1}$ | Curves derived on cut shoots | Rate of leaf water vapor loss |
| Leaf intrinsic water-use efficiency | WUEi | $\mu\text{mol mmol}^{-1}$ | Curves derived on cut shoots | Efficiency of water use during photosynthesis |
| Leaf optimal assimilation | $A_{opt}$ | $\mu\text{mol m}^{-2} \text{s}^{-1}$ | Curves derived on cut shoots | Maximum assimilation rate under optimal temperature conditions |
| Leaf temperature at optimal assimilation | $T_{opt}$ | °C | Curves derived on cut shoots | Temperature at which assimilation reaches its peak |
| Air temperature | $T_{air}$ | °C | Continuous point on *in-situ* shoots | Real-time air temperature measured directly in the field |
| Air maximum temperature | $T_{air,max}$ | °C | Meteorological-derived single point | Maximum daily air temperature derived from meteorological data |
| Leaf temperature | $T_{leaf}$ | °C | Continuous point on *in-situ* shoots | Real-time leaf temperature measured directly in the field |
| Leaf maximum temperature | $T_{leaf,max}$ | °C | Calculated single point | Maximum leaf temperature estimated from $T_{air}$ |
| Difference of leaf to air temperature | $\Delta T$ | °C | Calculated single point | Difference between $T_{leaf}$ and $T_{air}$ |
| Leaf thermal safety margin | TSM | °C | Calculated single point | Difference between $T_{crit}$ and $T_{leaf,max}$ |

Abbreviation: PSII: photosystem II.

### Environmental variables

Mean hourly minimum and maximum air temperatures, precipitation, humidity, wind speed, and solar radiation were recorded at a 10-minute temporal resolution by meteorological stations from the Meteo.cat network (Catalonia Meteorological Service). Gaps were filled with the *complete_meteo* function from the *meteoland* R package for the period of 2009 to 2024 (i.e., last 15 years). Approximately 0.05 % and 18 % of data points for humidity and solar radiation needed to be filled, respectively. Interpolation, using the *interpolator* function with the slope, elevation and aspect of each site was additionally conducted to downscale meteorological variables to each site (Table **1**). Average maximum monthly air temperature (MMAT) ranged between 18.78 and 27.82 °C for the coolest and the warmest site, respectively, over the last 15 years (from 2009 to 2024), and between 20.25 and 28.04 °C over the measurement period (2024) (Table **1**). The relationship between long *vs*. short-term MMAT across sites was highly significant (R² = 0.98 and p-value < 0.001; Fig. **S2**). Potential evapotranspiration was calculated at each site, using the Catalan Atlas for the period of 1951 to 1999, and following the Hargreaves and Samani (HS) model (Hargreaves & Samani, 1985). Precipitation divided by the potential evapotranspiration (P/PET) was then determined for each site from April to September as a measure of water availability. P/PET ranged between 0.22 and 0.91 for the driest and the wettest site, respectively, across the historical period from 1951 to 1999.

At each site, five soil samples (20 cm deep) evenly distributed over the total area were obtained using a soil core. The topsoil was discarded to exclude organic deposit and litterfall, and the five samples were combined for each site. The following variables were measured on each pooled sample for each site: organic carbon (OC; potentiometric titration), organic matter (OM; potentiometric titration), organic nitrogen (ON; volumetric titration), nitrate (NO_3_; UV-VIS spectrophotometry), phosphorus (UV-VIS spectrophotometry), clay, silt, and sand (sedimentation-Robinson pipette) at Eurofins Análisis Agro S.A. in Sidamon (Spain; Table **S3**).

To combine the different components of soil texture into a single, hydraulically meaningful variable, the coefficient *b* from the Saxton equation (Saxton *et al*., 1986), which governs the steepness of the soil water retention curve and is closely linked to hydraulic conductivity, was determined using the following equation:

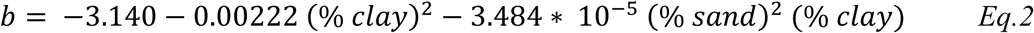

### Field measurement of leaf dark-adapted fluorescence and gas exchange temperature response curves

The photosynthetic efficiency of photosystem II (PSII; F_v_/F_m_) was measured at midday (between 14:00 and 15:00 local time) on three sunlight leaves per individual using a FluorPen FP100 (Photon Systems Instruments, Drásov, Czech Republic), reflecting maximum stress during midday depression. Leaves were covered with aluminum foil to maintain them into darkness for approximately 1 h before measurement, while 15 min is commonly accepted as a standard in the field (e.g., Giorio, 2011; Hara *et al*., 2021; Okubo *et al*., 2023a). Values were averaged within individuals. Measurements were obtained in a single day for all plants in each site.

Leaf gas exchanges were measured on one sunlight leaf per individual, except for *S. rosmarinus*, for which an average of 10 leaves was used to obtain sufficient leaf area, adjacent to those measured for F_v_/F_m_. Light-saturated photosynthesis (A), stomatal conductance (g_s_) and transpiration (E) rates were measured in the field at different air temperatures to build temperature response curves using a LI-COR 6400XT infrared gas analyzer system (LI-COR, Lincold, USA). The broadleaf chamber (2 cm²) was used for all the individuals, including *S. rosmarinus*, to ensure measurement reliability and comparability across species. The leaf enclosed in the cuvette was harvested if it was smaller than the chamber, and the projected leaf area, which was used to correct recorded values, was measured using a flatbed scanner (Epson Perfection V39 II, Amsterdam, Netherlands). One meter-long branches were collected and placed in a water bucket and recut twice to avoid potential cavitation (Bachofen *et al*., 2020). Although the cut-branch approach is commonly used to measure gas exchange (Santiago & Mulkey, 2003; Miyazawa *et al*., 2011), it should be noted that these rates may be lower than pre-excision ones. Nevertheless, studies have shown that bias is minimized when the length of the excised branch exceeds the xylem vessel (Missik *et al*., 2021). Therefore, while some underestimation of gas exchange values is possible, it is likely to be minimal. Measurements were carried out from sunrise (between 07:30 to 08:00 local time) until the peak of daily temperatures (between 13:00 and 14:00 local time) following the natural daily variation of ambient air temperatures. The air temperature inside the cuvette was set to match the ambient air temperature, tracking the ambient air temperature fluctuation while minimizing artefacts caused by temperature differences between the natural environment and the cuvette conditions. An additional measurement at ambient air temperature + 5 °C was taken at the end of the day to obtain additional extreme conditions, leading to an optimal total of seven measurements across a temperature gradient per individual and day. Measurements were obtained over one to three consecutive days for each site (depending on the individual number per site). To prevent confounding factors in the interpretation of temperature curves, the following settings were fixed: 400 ppm of reference CO_2_ concentration, 1500 μmol m⁻² s⁻¹ light-saturating photosynthetic photon flux density, and relative air humidity set at maximum to maintain VPD as low as (but reaching up to 9 kPa at the highest temperatures; Fig. **S3**). Although this method still highly confounds temperature with VPD effects, it still provides useful relevant field-insights. Measurements were recorded after steady-state gas exchange rates were maintained for at least 3 min. Intrinsic water-use efficiency (WUE_i_) was calculated as described in Fischer & Turner (1978):

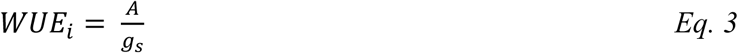

Photosynthesis (A) was plotted against the leaf temperature measured by the LI-COR chamber (T_leaf, LI-COR_) (i.e., A *vs*. T_leaf, LI-COR_; Fig. **S4**) and fitted with a non-linear regression following a second order Gaussian function as described in June *et al*. (2004) and Slot & Winter (2017):

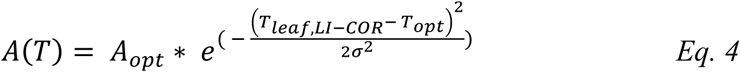

where A(T) represents the light-saturated photosynthesis at a given T_leaf, LICOR_, σ describes the spread of the temperature response curve, and A_opt_ is highest assimilation rate occurring at the optimal temperature (T_opt_; Fig. **S4**).

### Field monitoring of leaf and air-temperature

Leaf and air temperature (T_leaf_ and T_air_, respectively) were recorded on one sunlight leaf adjacent to those measured for F_v_/F_m_ and leaf gas exchanges temperature response curves. Sensors were always positioned in the same orientation on the upper leaf surface, avoiding the major vein. These variables were measured for six of the selected individuals per species at each site using double-ended LAT-B3 sensors (Ecomatik, Germany) connected to two CR1000 datalogger (Campbell Scientific, UK). Data was recorded and stored every 10 min for a total period of at least 24h per individual (between 14:00 and 18:00 local time the following day). These sensors are specifically designed for field measurements up to +70°C, with a tolerance for the leaf-to-air temperature difference of approximately ±0.2°C across the full temperature range. Additionally, any unrealistic values were removed when necessary. The difference between leaf and air- temperature (T_leaf_ – T_air_ = ΔT) was calculated for each individual.

### Laboratory measurement of leaf thermotolerance curves and thermal thresholds

The thermotolerance curves were performed on one excised sunlight leaf per individual adjacent to those measured for F_v_/F_m_, leaf gas exchange temperature response curves, T_leaf_ and T_air_. Leaves were harvested at midday (between 14:00 and 15:00 local time), placed in a sealed plastic bag, and stored in a dark and cool room (approx. 20 °C) for a maximum of 20 h until the next days’ analysis. Even if some recent studies showed that waiting time until measurement does not significantly affect leaf thermotolerance curves (Tarvainen *et al*., 2022), we acknowledge that prolonged storage may still introduce minor artifacts. The thermotolerance curves were obtained by submerging the leaves, hermetically sealed into transparent plastic bags, into a water bath (Ovan, Spain) under rising temperatures at 5 °C intervals from 30 to 65 °C (i.e., 30, 35, 40, 45, 50, 55, 60, 65 °C). The same leaves were exposed to each temperature, simulating the progressive heat that occurs under natural environments (e.g., as acknowledge in recent dose-response experiments; Neuner & Buchner, 2023). Leaves were kept at each temperature for 15 min as this exposure is sufficiently long to stabilize physiological processes (see Marias *et al*., 2017; Gauthey *et al*., 2023; Didion-Gency *et al*., 2025). The bath temperature was continuously monitored with an analogue thermometer inserted into the water. The stepwise temperature increase was applied using a single water bath and the water bath temperature was changed between temperature steps. After exposure to a given temperature, plastic bags were immediately dried and stored in the dark for 15 min (following e.g., Okubo, Inoue & Ishii, 2023). Afterwards, F_v_/F_m_ was measured using a FluorPen FP100 (Photon Systems Instruments, Drásov, Czech Republic). The thermotolerance curves (i.e., F_v_/F_m_ *vs*. bath temperature; Fig. **S5**) were plotted and fitted with separate third-order sigmoid functions as described in Marias *et al*. (2016). Critical leaf temperature (T_crit_) was extracted as the intersection point between the linear regression fitted to the high, slow changing segment and the rapidly declining segment of F_v_/F_m_ (Schreiber & Berry, 1977; Knight & Ackerly, 2002). The maximum temperature (T_max_) was extracted as the intersection point between the rapidly declining segment and the low, slowly changing segment of F_v_/F_m_ (Didion-Gency *et al*., 2025). The temperature causing a 50% reduction of F_v_/F_m_ (T_50_) was extracted as the midpoint between T_crit_ and T_max_ (Marias *et al*. 2017). A sigmoidal Weibull model was used to generate the final figures, as it provides a continuous curve that incorporates all data points and enhances the visualization of both thresholds (Fig. **S5**). Thermal thresholds (T_crit_, T_50_, T_max_), derived from the raw data, are also shown for reference (Fig. **S5**).

The thermal safety margin (TSM) was calculated as follows for each individual:

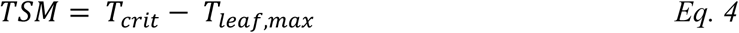

where T_leaf,max_ represents the hourly maximum leaf temperature, determined using T_air,max_ from the interpolated data from Meteo.cat 2024 (Table **1**), corrected for the specific relationship between T_air_ and measured T_leaf_ for each species (Fig. **S6**).

### Statistical analysis

All analyses were performed using the R v.4.5.2 statistical platform (Rstudio, 2025). All traits were visually inspected for normality and homoscedasticity, both on the raw data and on model residuals, and were also checked for normality using Shapiro-Wilk test (*shapiro.test* function; Shapiro & Wilk, 1965). Log-transformations were only applied when these inspections indicated it was necessary.

A preliminary analysis was conducted to assess the impact of repeated gas exchange measurements on individuals from PAS, as this site was visited twice: once at the beginning of the measurement period (where only A_opt_ and T_opt_ from three individuals could be extracted with a minimum of five points per individual due to equipment failure) and once at the end. Differences between the two periods (for the same individuals) were tested using a paired t-test (*t.test* function; Gayen, 1949), resulting in a *p-value* = 0.056 and *p-value* = 0.181, for A_opt_ and T_opt_, respectively. However, only A_opt_ and T_opt_ of three individuals between the first and second repetition could be compared (n = 3 individuals), implying that the results for A_opt_ and T_opt_ should still be interpreted with caution for the PAS site.

An exploratory analysis on soil variables was performed using a Principal Component Analysis (PCA), with variable standardization and self-adjust axis rotation (*prcomp* function; Holland, 2019). PCA was used to investigate and reveal the structures of variability and correlation between the soil characteristics of the study sites (i.e., OC, OM, ON, NO_3_, clay, silt, sand and *b*). Phosphorus was removed from this analysis as the determination was not precise enough to get exact values (i.e., concentration always shown below 5% for all sites; Table **S3**). *b* (13.06 contribution to PC2, Table **S4**) was selected for the following statistical analysis due to its strong association with water retention and texture (Fig. **S7**).

Similarly, an exploratory analysis combining the environmental and shrubs variables was performed using a PCA, with variable standardization and self-adjust axis rotation (*prcomp* function; Holland, 2019). We included the following environmental and shrubs variables of the species and study sites: MMAT 2024, MMAT 2009 – 2024, P/PET, H, BD, SN, P_competition_, P_crown_, *b*, as well as the first axis scores of the soil PCA. MMAT 2024, H and P_competition_ (0.86, 2.98, and 0.05 contribution to PC1, PC2 and PC2, respectively, Table **S5**) were selected because they are relatively independent descriptors of climate, plant size and its competitive environment, respectively (Fig. **S8**).

The responses of F_v_/F_m_, T_crit_, T_50_, T_max_, A_opt_, T_opt_ and TSM to the additive effects of MMAT 2024, P_competition_, H, and *b* were determined through linear and quadratic mixed-effects models (*lme4* and *car* functions; Fox *et al*., 2013; Bates, 2018). The quadratic mixed-effects model was selected when it significantly improved model fit, as indicated by a lower Akaike Information Criterion (AIC; Akaike, 1998). In the absence of such improvement, the simpler linear mixed-effects model was chosen in. The individual site and species were treated as random effects to account for variation among sites and among species (but see species- and site-level random estimates, Figs. **S9**, **S10** & **S11**, as well as the corresponding species- and site-specific values, Fig. **S12**). P/PET was not included in the models as it is a confounding factor of MMAT 2024 (Fig. **S8**), and the focus of our study is on temperature response.

Similarly, the responses of T_leaf_ and ΔT to the interactive effect of T_air_ by species (i.e., Ao, Au, Bs, Pl, Ra, and Sr), and the additive effect of P_competition_, H, and *b* were determined through linear mixed-effects models (*lme4* and *car* functions; Fox *et al*., 2013; Bates, 2018), where the individual site was treated as random effect. Additionally, the relationship between ΔT and T_air_ was examined using linear models (*lm* functions) for each species across sites, but no significant differences in slope were found in any species (Table **S6**), supporting the use of species-level-only shown relationship.

The overall responses of A to the interactive effects of g_s_ and T_leaf_, and E and T_leaf_, as well as the additive effects of P_competition_, H, and *b* were determined through linear mixed-effects models (*lme4* and *car* functions; Fox *et al*., 2013; Bates, 2018). However, as *in-situ* T_air_ and VPD_air_ inside the LI-COR chamber were highly correlated (Fig. **S3**), we also fitted an alternative model that replaced T_leaf_ with leaf VPD (VPD_leaf_) in the interactions, while retaining the same additive terms. The T_leaf_ mixed-effects models was chosen as VPD_leaf_ did not significantly improved model fit, as indicated by a similar or higher AIC (Akaike, 1998; Tables **S7** & **S8**). However, to still account for the potential residual effect of VPD_leaf_, we orthogonalized VPD_leaf_ by regressing it on T_leaf_ and include the resulting residuals (VPD_leaf,resid_) in the final model linear mixed-effects models. The 5^th^ and 95^th^ percentiles of T_leaf_ were used to illustrate the interactive effect of T_leaf_ on the relationships between A and g_s_ or E. The individual site and species were treated as crossed random effects.

## Results

### Relationship between leaf thermal tolerance traits and warmer temperatures

According to our models, the air temperature gradient across sites significantly impacted four traits (i.e., leaf photosynthetic efficiency of the photosystem II, F_v_/F_m_, leaf maximum temperature, T_max_, leaf optimal temperature, T_opt_, and leaf thermal safety margin, TSM) related to leaf thermal tolerance across species. F_v_/F_m_ and T_max_ showed a quadratic relationship with monthly maximum air temperature (MMAT), with an initial increase and a decline beyond a certain MMAT threshold (i.e., 24.1 °C and 24.5 °C , respectively, Fig. **2**, Table **S9**). In contrast, T_opt_ significantly increased and TSM significantly decreased in response to higher MMAT (Fig. **2**, Table **S9**). No effect of MMAT was found on the leaf critical temperature (T_crit_), temperature causing a 50 % reduction of F_v_/F_m_ (T_50_), and optimal assimilation (A_opt_; Figs. **2** & **S9**, Table **S9**). On average T_crit_ and T_50_ were approx. 42.2 ± 2.2 °C and 50.8 ± 2.2 °C, respectively, and A_opt_ was approx. 2.9 ± 2.9 µmol m^-2^ s^-1^.

**Figure 2:**
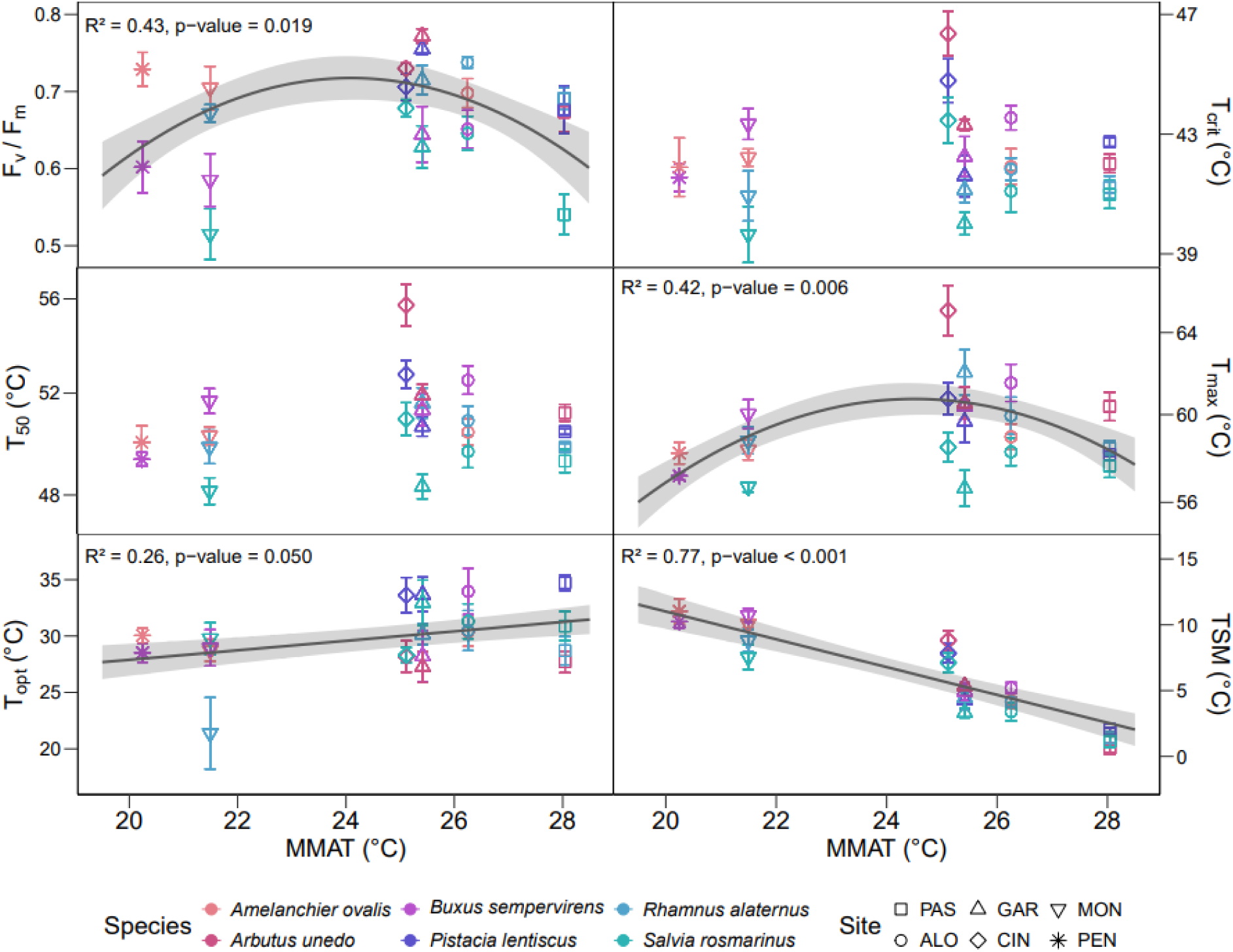
Photosynthetic efficiency of the photosystem II (F_v_/F_m_), critical leaf temperature (T_crit_), temperature causing a 50% reduction of F_v_/F_m_ (T_50_), maximum temperature (T_max_), leaf optimal temperature (T_opt_), and leaf thermal safety margin (TSM) as a function of the maximum monthly air temperature (MMAT) for 2024 for each species (i.e., *A. ovalis*, *A. unedo*, *B. sempervirens*, *P. lentiscus*, *R. alaternus*, and *S. rosmarinus*) and sites (i.e., PAS: Pas de l’Ase, ALO: Alòs de Balaguer, GAR: Garraf, CIN: Cingles de Bertí, MON: Montsec, PEN: Pentina; mean ± SE, *n* = 8 shrubs per site and species). Significant regression lines are shown for the corresponding mixed effects models when the overall model was significant (*p*-values ≤ 0.05; mean ± SE). Variables were log-transformed whenever required to satisfy normality assumptions. The R² indicates and p-value indicate significance of the relationship.

### Relationship between leaf and air temperature

The relationship between air temperature (T_air_) and leaf temperature (T_leaf_), as well as the relationship with the difference of leaf and air temperature (ΔT), were significant for all species. All species showed significantly higher T_leaf_ and ΔT with increasing T_air_ (Figs. **3** & **S6**, Table **S10**). Specifically, T_leaf_ increased between 1.09 and 1.24 °C for every 1 °C increase in T_air_ across all species (Fig. **S6**). Similarly, ΔT increased between 0.09 and 0.24 °C for every 1 °C increase in T_air_ across all species (Fig. **3**). Moreover, while ΔT showed mostly values above 0 during the daytime (i.e., 9:00 to 21:00), and rarely dropped below 0, we still observed substantially negative daytime values, particularly for *R. alaternus* and *S. rosmarinus* and, to a lower extent, *A. ovalis* (Fig. **3**).

**Figure 3:**
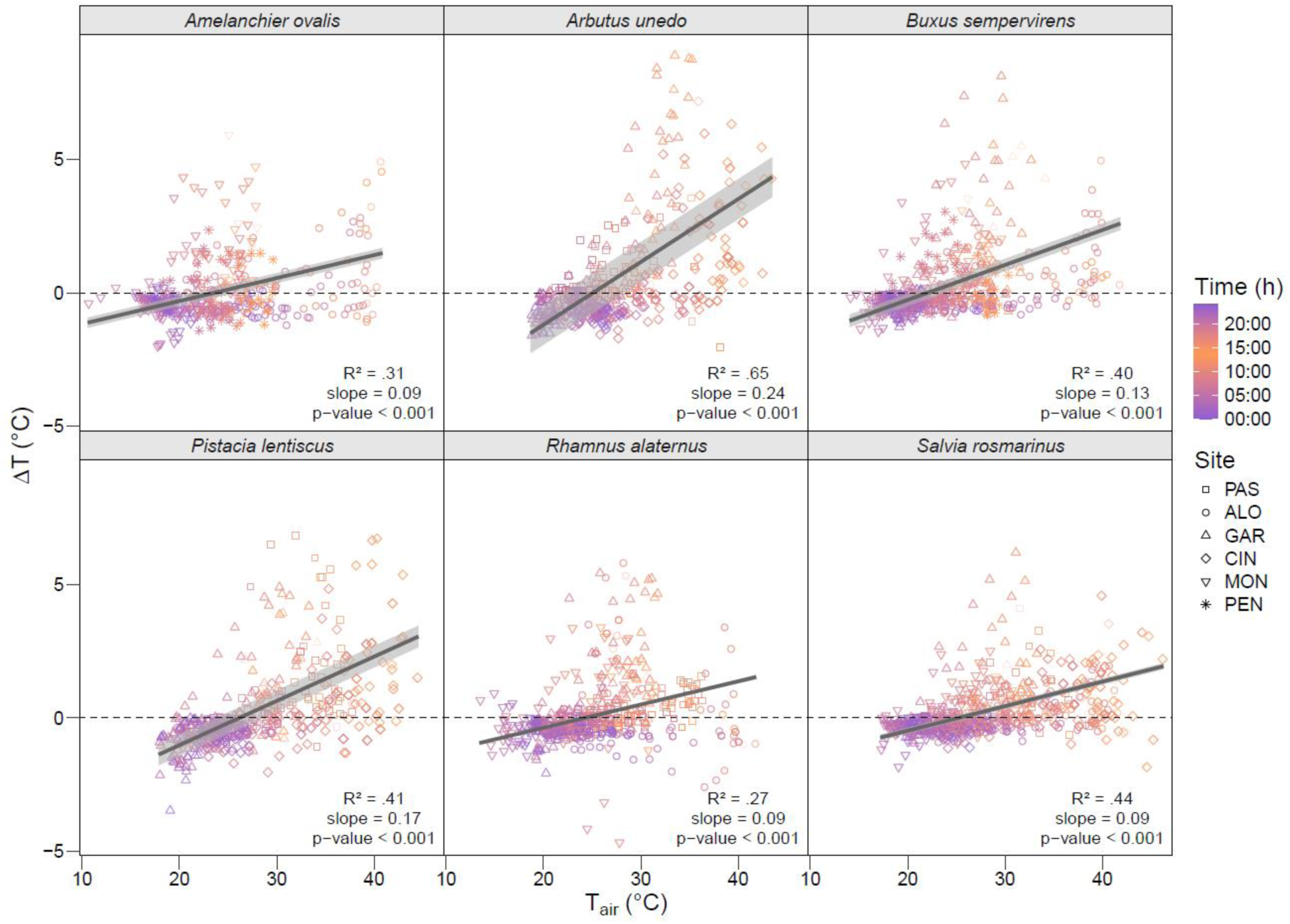
Relationship between the difference of leaf and air temperature (ΔT) and the air temperature (T_air_) for each species (i.e., *A. ovalis*, *A. unedo*, *B. sempervirens*, *P. lentiscus*, *R. alaternus*, and *S. rosmarinus*) and sites (i.e., PAS: Pas de l’Ase, ALO: Alòs de Balaguer, GAR: Garraf, CIN: Cingles de Bertí, MON: Montsec, PEN: Pentina). Color gradients represent the hours of the day. The dashed line represents when there is no difference between T_leaf_ and T_air_, while the solid grey line represents the linear relationship between ΔT and T_air_ for the corresponding mixed effects models. The R² indicates, slope and p-value indicate significance of the relationship.

### Relationship between leaf gas exchanges and leaf temperature

Across sites and species, the relationships between leaf assimilation (A) and stomatal conductance (g_s_), and transpiration (E) were positive and highly significant (Fig. **4**, Table **S11**).

**Figure 4:**
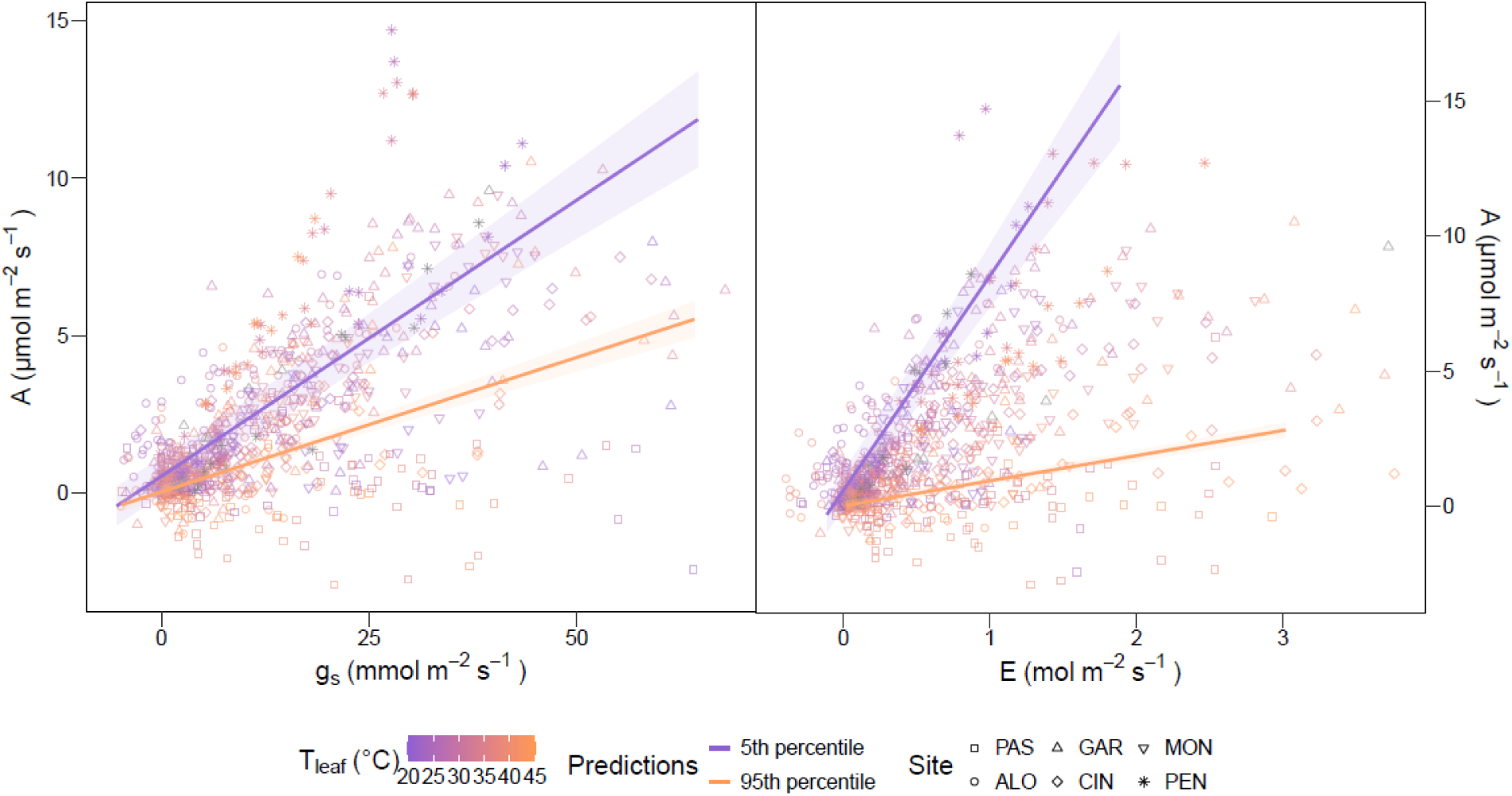
Relationship between leaf assimilation (A) and leaf stomatal conductance (g_s_) and leaf transpiration (E) across all species (i.e., *A. ovalis*, *A. unedo*, *B. sempervirens*, *P. lentiscus*, *R. alaternus*, and *S. rosmarinus*) and sites (i.e., PAS: Pas de l’Ase, ALO: Alòs de Balaguer, GAR: Garraf, CIN: Cingles de Bertí, MON: Montsec, PEN: Pentina). Color gradients represent the leaf temperature (T_leaf_). To illustrate the interaction between g_s_ or E and T_leaf_, we show the linear regression lines for the 5^th^ and 95^th^ percentile of T_leaf_ (purple and orange, respectively).

Both g_s_ and E were positively related to A, and the slope depended on leaf temperature, as indicated by the significant interaction between g_s_ (or E) and T_leaf_ (Table **S11**). Specifically, the slope of the relationship between A and g_s_ or E weakened at higher T_leaf_, as graphically shown by contrasts the relationship for the 5^th^ and 95^th^ percentiles of T_leaf_ (Fig. **4**, Table **S11**). Additionally, leaf VPD residuals (VPD_leaf,resid_) exhibited a significant negative relationship with A, indicating that elevated leaf-level evaporative demand tended to reduce A (Table **S11**). Moreover, A exhibited an overall quadratic relationship with rising T_leaf_, with an initial increase followed by a decline beyond a certain threshold (Fig. **S4**), which allowed us to identify A_opt_ and T_opt_. At the highest T_leaf_ measured (40-45 °C), A dropped to levels comparable to those observed at around 20 °C (Fig. **S4**). In contrast, E exhibited increases with rising T_leaf_ in most sites and species (Fig. **S13**), whereas the relationship between g_s_ and T_leaf_ was more variable but still showed mostly stable or parabolic trends depending on the site (Fig. **S14**). Additionally, leaf intrinsic water-use efficiency (WUE_i_) remained mostly stable with rising T_leaf_ in most sites (Fig. **S15**).

## Discussion

### Impact of warmer temperatures on leaf thermal tolerance traits

As we initially expected, our study demonstrated that Mediterranean shrubs have some capacity to adjust warmer conditions (i.e., higher leaf optimal temperature, T_opt_; Fig. **2**). Our results agree with the scarce studies investigating heat impacts on optimal temperature of photosynthesis reporting an increase in T_opt_ for plants growing in warmer climates (Berry & Bjorkman, 1980; Cabrera *et al*., 1998; Yamori *et al*., 2014; Deluigi *et al*., 2025). However, most of our measured traits, either did not change (i.e., leaf critical temperature, T_crit_, and temperature causing a 50 % reduction of F_v_/F_m_, T_50_) or showed a quadratic relationship (i.e., leaf photosynthetic efficiency of the photosystem II, F_v_/F_m_, and leaf maximum temperature, T_max_) with monthly maximum air temperature (MMAT). These quadratic relationships implied that, beyond a given threshold, F_v_/F_m_ and T_max_ tended to decrease with temperature. As a result, the leaf thermal safety margin (TSM) showed a negative relationship with MMAT and got close to 0 at the warmest sites (Fig. **2**). Thus, our results suggest that the adjustments in leaf thermal tolerance traits do not compensate for the increased exposure at the warmer sites, and therefore shrubs are operating closer to their thermal limits. These findings support recent studies indicating that such leaf thermal adjustments are often insufficient and that prolonged exposure to warmer climates will inevitably cause irreversible damage (O’sullivan *et al*., 2017; Zhu *et al*., 2018). This highlights the importance of incorporating thermal vulnerability alongside drought when predicting vegetation dynamics in Mediterranean communities.

Additionally, while we did not observe a significant impact of warmer climates on T_crit_ and T_50_ (Fig. **2**), previous studies in tropical species have shown considerable variation in these traits across sites and species (Sastry & Barua, 2017; Slot *et al*., 2019; Perez *et al*., 2021), though this variation may not necessarily reflect direct adjustments to thermal conditions. T_crit_ may indicate the initial stage of damage to F_v_/F_m_ (Maxwell & Johnson, 2000b), which, on average, occurred at 42.2 ± 2.2 °C. At lower temperatures (T < T_crit_), heat-induced denaturation of lipids and proteins in the thylakoid membrane is unlikely to have accumulated yet, as heat shock proteins may help counteract early thermal stress (Thebud & Santarius, 1982; Yordanov *et al*., 1986; Eaton-Rye *et al*., 2012). However, once temperatures exceed T_crit_ (T ≥ T_crit_), heat stress can impair leaf enzymatic functions, such as Rubisco (Bose *et al*., 1999; Bernacchi *et al*., 2002; Salvucci & Crafts-Brandner, 2004). An important consideration is that our study focused on Mediterranean shrubs, suggesting that variation in the earliest thermal thresholds (T_crit_ and T_50_) might be related to species type or the growing habitats (i.e., tropical *vs*. Mediterranean). Similarly, Gauthey *et al*. (2023) did not observe any adjustment of T_crit_ and T_50_, determined using the minimal chlorophyll-*a* fluorescence, F_0_, in Mediterranean coniferous tree species. Furthermore, no genetic-based assessments were conducted on the selected individuals along the climatic gradient. Thus, the lack of adjustment could also reflect a limited genetic differentiation among populations, potentially constraining T_crit_ and T_50_. Moreover, our experimental approach primarily captures long-term adjustment, yet tolerance to temperatures may also be expressed through short-term plasticity in these plastic traits, which could additionally contribute to the observed patterns. Additionally, leaves were held overnight at 20°C prior measurement, which could potentially reduce some natural variation in the earliest thermal thresholds (i.e., T_crit_ and T_50;_ Drake *et al*., 2018; Neuner & Buchner, 2023). Nevertheless, waiting time showed to not significantly affect leaf thermotolerance curves (Tarvainen *et al*., 2022), so any artifacts are likely minor. Therefore, both limited genetic differentiation and short-term plastic responses remain plausible explanations, and further work disentangling genetic *vs*. environmental effect on intraspecific variability would be needed to better understand these mechanisms. Nonetheless, understanding these adjustments under natural conditions remains an important research priority.

### Leaf temperature regulation and gas exchange decoupling

Our results demonstrate that an increase in air temperature (T_air_) led to positive differences between leaf temperature (T_leaf_) and T_air_ (ΔT), with a severe overheating up to 8 °C above T_air_ for *A. unedo* (Fig. **3**), the species with largest leaves. Such positive ΔT values do not arise solely from rising T_air_, but also because leaves absorb more solar radiation than the surrounding air in sunny conditions (Sirvydas *et al*., 2010). These findings support other studies that observed T_leaf_ differing from T_air_, with some cases where T_leaf_ can exceed T_air_ up to 20 °C (Salisbury & Spomer, 1964; Doughty & Goulden, 2008). Similarly, some T_leaf_ were also found to be lower than T_air_, with the most negative ΔT occurring primarily during night, early morning or evening times (Fig. **3**). This indicates that leaves are cooler than T_air_ during low radiation periods (Rey-Sánchez *et al*., 2016; Miller *et al*., 2021), due to their higher thermal conductivity in darkness.

Nevertheless, negative ΔT values at high temperatures were also observed during periods of high radiation (i.e., daytime) for some species, such as *R. alaternus* and *S. rosmarinus* (Fig. **3**). In these cases, the excess energy input from strong solar radiation may have been dissipated through a passive or active evaporative cooling, achieved by maintaining relatively high stomatal conductance and transpiration (Schymanski *et al*., 2013; Marchin *et al*., 2016; Urban *et al*., 2017; Drake *et al*., 2018a), and/or a continued water loss through stomatal and cuticular leakage (Duursma *et al*., 2019; Fernandes *et al*., 2025). Additionally, leaf inclination may also contribute to reduce radiation absorption, as steeper (i.e., more vertical) leaf angles tend to intercept less radiation, and therefore experience lower heat load compared to more horizontal ones (Vogel, 1970; Zhou *et al*., 2023). Nonetheless, it should be noted that leaf inclination was not measured in this study, and thus its potential contribution remains unquantified.

Yet, several recent studies have shown evidence of a decoupling between leaf assimilation (A) and stomatal conductance (g_s_) or transpiration (E) at high temperatures (Marchin *et al*., 2023; Diao *et al*., 2024; Gauthey *et al*., 2024), with potential major implications for carbon and water relations, particularly under drought. Our study showed similar results, where the relationship between A and g_s_ or E depended on T_leaf_, with weaker slopes under higher temperature (e.g., 95^th^ *vs*. 5^th^ percentile of T_leaf_; Fig. **4**). These results indicate a partial shift towards decoupling of gas exchange in Mediterranean shrubs at high temperatures, with a significant reduction in carbon uptake efficiency without its corresponding decline in water loss (Fig. **4**). Gas exchange decoupling was first expected to occur only under well-watered conditions, as prolonged water loss through evaporating cooling could lead to xylem embolism and hydraulic failure (Urban *et al*., 2017; Drake *et al*., 2018a). However, recent work showed that gas exchange decoupling was also possible under soil drought conditions and when VPD reached extreme values (Marchin *et al*., 2023). Similarly, our results revealed that VPD residuals (VPD_leaf,resid_) had a significant negative effect on A, independently of g_s_, E, or T_leaf_ (Table **S11**), indicating that increases in evaporative demand can directly suppress carbon assimilation even when stomatal responses alone do not predict such declines. Rising VPD may therefore amplify gas exchange decoupling by imposing additional physiological constraints beyond those caused by temperature alone. Thus, together, our results expand these recent findings and show evidence of gas exchange decoupling even in drought-prone Mediterranean shrubs at the peak of the summer, when soil moisture is typically low. A limitation of our work was that our experimental approach mixes the effects of different climatic (i.e., temperature, VPD and soil drought) and topo-climatic (i.e., elevation) drivers. While this approach does not allow us to fully disentangle the mechanisms underlying gas exchange decoupling, our results support the idea that this decoupling may occur more frequently than previously recognized, including under natural conditions in Mediterranean shrubs.

Furthermore, although the mechanisms driving gas exchange decoupling remain poorly understood, an active evaporative cooling or passive water loss through stomatal and cuticular leakage have been suggested as potential contributors. Our results showed that gₛ was mostly stable, while E consistently increased as T_leaf_ raised (Figs. **S13** & **S14**), suggesting that stomatal closure likely occurred to reduce water loss, while a passive water loss through stomatal leakage and cuticular conductance might arise. A comparison with the leaf residual conductance (g_res_), measured at 25°C on the same individuals using the gravimetric method (data not shown; Method **S1**), showed that g_s_ mostly did not exceed g_res_ at the highest measured T_leaf_ (above 40°C), except for *A. ovalis* and *A. unedo* (Fig. **S16**). These results indicate a nearly-complete stomatal closure for most species, and are hence suggestive of passive cooling. Interestingly, *A. ovalis* and *A. unedo* had the largest studied leaves, suggesting that leaf morphological traits, such as greater leaf area often associated with more extensive veinlet networks (Zhu *et al*., 2012), or sparse but large stomata (Leuschner & Ellenberg, 2017), could support an active evaporative cooling in these two species. Yet, to our knowledge, no study has examined the direct influence of morphological traits on evaporative cooling, highlighting an important direction for future research. However, this comparison (g_s_ *vs*. g_res_) should still be interpreted with caution, as here g_res_ was measured at 25°C and assumed to be constant across temperature, likely underestimating the actual g_res_ at higher leaf temperatures. Future work should include a full temperature-response curve of g_res_ using the gravimetric technic to more accurately constrain the g_s_ *vs.* g_res_ comparison. Nonetheless, the increase in E with rising T_leaf_ could be attributed to both a higher evaporative demand driven by increased VPD and a reduction in water viscosity to facilitate water transport, which has been shown to improve the water supply to evaporative sites within the leaf intercellular cavities (Fredeen & Sage, 1999; Diao *et al*., 2024). Similarly, Cochard *et al*. (2000) also showed that increased water fluidity enhances whole-plant hydraulic conductance, thereby improving water uptake and supporting higher transpiration rates without lowering leaf water potential, enhancing longer-term survival under high temperature.

Additionally, our results also showed an intrinsic water-use efficiency (WUE_i_) approaching zero across all temperatures (Fig. **S15**). Low temperatures (T < 25 °C) reduced both A and g_s_, moderate temperatures (25°C < T < 35 °C) maintained high A alongside high g_s_, and high temperatures (T > 35 °C) caused A to collapse while g_s_ remained positive (Figs. **S4** & **S14**), leading to near-zero WUE_i_ across the full temperature range, and implying the presence of non-stomatal limitations to photosynthesis at high temperatures. Previous research has shown that Rubisco deactivation or degradation could occur after an exposition of 4 min at 35 °C (Salvucci & Crafts-Brandner, 2004b), a temperature similar to that experienced by our study plants when A began to decrease (Fig. **S4**). Similarly, mesophyll conductance has also been shown to limit photosynthesis under warm and dry conditions (Bernacchi *et al*., 2002; Urban *et al*., 2017; Sperlich *et al*., 2019), partly due to the downregulation of aquaporins which impairs internal CO_2_ diffusion within the leaf (Uehlein *et al*., 2008; Chen *et al*., 2023), and in some case was shown to be greater than stomatal limitation (Zhu *et al*., 2021). It is also known that mesophyll conductance tends to increase with temperature also in the absence of drought (Von Caemmerer & Evans, 2015), so further work would be needed to fully explore the mechanisms behind these responses.

Although gas exchange decoupling through the maintenance of E at high temperatures might not be sufficient to substantially lower T_leaf_, it might help prevent it from reaching detrimental levels. Indeed, despite extremely high T_air_ during the summer (Tables **1** & **S2**), our results showed that only two individuals at the warmer site entered the thermal danger zone (TSM < 0) during the study period, while the remaining individuals maintained positive TSM (Fig. **5a**). However, in the absence of T_crit_ adjustment, and given the projected increases in the frequency and intensity of extreme heatwaves (Park Williams *et al*., 2013; IPCC, 2023), TSM might be exceeded to narrow in the coming decades. For instance, the cumulative number of days during which estimated T_leaf_ exceeded T_crit_ over the last 15 years reached up to 294 days for our *S. rosmarinus* and > 100 days in *A. unedo* and *R. alaternus* across all sites combined, with the warmest site being the most affected (i.e., PAS; Fig. **5b**). This could result in widespread leaf scorching and tissue damage in Mediterranean shrubs and potentially accelerate vegetation mortality at regional scales (Breshears *et al*., 2021; Gazol & Camarero, 2022; Gong & Hao, 2023). Importantly, although our TSM values are based on single-point measurements of T_crit_, and therefore do not capture potential short-term or seasonal adjustments, our measurements were collected when each site was consistently near its seasonal thermal maximum (Table **S2**), enhancing the robustness of our thermal-vulnerability assessments across sites and species. However, a limitation of our TSM is that our T_crit_ was estimated using 15-min exposures, whereas our T_leaf,max_ was derived from 10-min observations. Recent studies showed that exposure duration has a strong influence on thermal stress (Neuner & Buchner, 2023; Faber *et al*., 2024; Cook *et al*., 2024; Arnold *et al*., 2025), leading to a possible slight overestimation. Nevertheless, to our knowledge, most vegetation models predicting leaf carbon and water balance assume a linear relationship between gas exchange variables (e.g., Harley & Baldocchi, 1995). With these recent findings, models may become inaccurate at high temperatures, highlighting the urgent need to further develop vegetation models that can account for the full range of responses to extreme temperatures.

**Figure 5:**
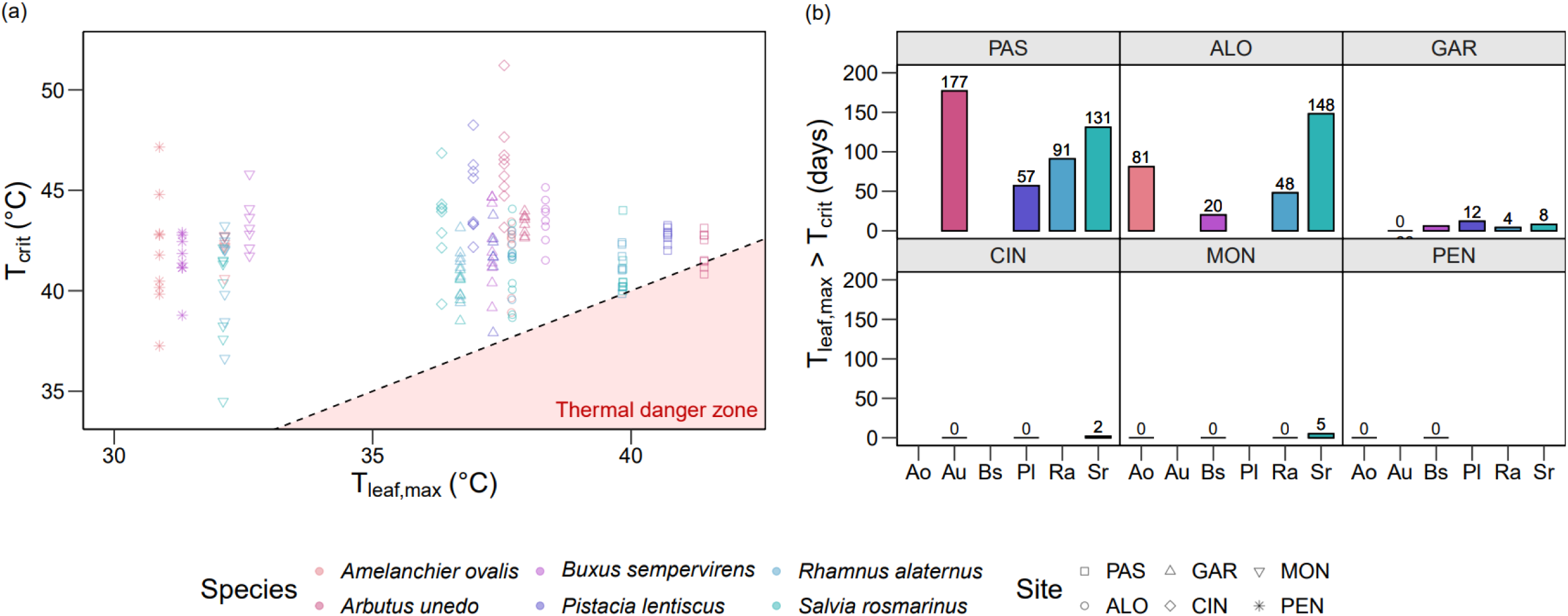
(a) Relationship between critical leaf temperature (T_crit_) and maximum leaf temperature experienced in 2024 (T_leaf,max_) across all species (i.e., *A. ovalis*, *A. unedo*, *B. sempervirens*, *P. lentiscus*, *R. alaternus*, and *S. rosmarinus*) and sites (i.e., PAS: Pas de l’Ase, ALO: Alòs de Balaguer, GAR: Garraf, CIN: Cingles de Bertí, MON: Montsec, PEN: Pentina). Each point represents an individual measurement. The dashed line marks the threshold where T_leaf,max_ equals T_crit_, and the shaded area below this line indicates the thermal danger zone, where T_leaf,max_ exceeds the plant’s thermal tolerance threshold. (b) Cumulative number of days during which T_leaf,max_ exceeded T_crit_ over the last 15 years (from 2009 to 2024) for each species and sites.

## Conclusion

Our study highlights that Mediterranean shrubs show limited capacity to adjust their leaf thermal tolerance traits across a climatic gradient, resulting in narrower TSM in warmer sites. These results suggest that shrubs may be operating close to their thermal limit as conditions get warmer, indicating a high vulnerability to rising air temperatures and more frequent and intense heatwaves projected under global warming. Our work further revealed that T_leaf_ could be lower than T_air_ at high temperatures due to evaporative cooling processes. This cooling effect might be maintained by passive water loss through stomatal and cuticular leakage, which supports relatively high transpiration rates at high temperatures. These processes, together with non-stomatal limitations to photosynthesis under high temperatures, have led to an observation of widespread gas exchange decoupling in our Mediterranean shrubs, suggesting that this mechanism may occur more frequently than previously recognized, especially in drier climates with high VPD. Even if thermal safety margins were mostly not exceeded during the study period, our study suggests that leaf thermal adjustments and evaporative cooling processes may become insufficient in the future, and projected prolonged exposure to extreme heat is likely to result in irreversible leaf damage. These findings also highlight the need to account for direct heat effects and gas exchange decoupling in future vegetation dynamics models for Mediterranean shrubs.

## Acknowledgements and funding

MD-G was supported by the competitive Post.doc mobility grant P500PB_222186 funded by the Swiss National Science Foundation (SNSF). PS was supported by the competitive grant PRE2022-103498 funded by the Agencia Estatal de Investigación (MCIN/AEI/10.13039/501100011033). This research was supported by the competitive grant PID2021-127452NB-I00 (TRACES) funded by the Agencia Estatal de Investigación (MCIN/AEI/10.13039/501100011033). JM-V acknowledges the Agència de Gestió d’Ajuts Universitaris i de Recerca de Catalunya (grant 2021 SGR 00849) and the Institució Catalana de Recerca i Estudis Avançats (ICREA Academia award). We thank S. Palau-Saez, A. Sanmartin- Arevalo and J. Fernandez-Boixaderas for their help during fieldwork. We are grateful to Á. Vàzquez-Làzaro for his contribution to data acquisition in the laboratory.

## Author contributions

All authors conceived and designed the study together; MD-G and PS collected the data; MD-G analyzed the data and led the writing of the manuscript. All authors critically contributed to the manuscript and gave final approval for publication.

## Supplementary information

Supplementary information is available for this paper.

## Data availability statement

Data used in this manuscript will be available from the Dryad Digital Repository after acceptation of the manuscript. Data supporting the findings of this study will also be available from the corresponding author, MD-G.

## References

1. Ahrens CW, Challis A, Byrne M, Leigh A, Nicotra AB, Tissue D, Rymer P. 2021. Repeated extreme heatwaves result in higher leaf thermal tolerances and greater safety margins. New Phytologist 232: 1212–1225.

2. Akaike H. 1998. Information Theory and an Extension of the Maximum Likelihood Principle. Breakthroughs in Statistics I: 310–324.

3. Allen CD, Breshears DD, McDowell NG. 2015. On underestimation of global vulnerability to tree mortality and forest die-off from hotter drought in the Anthropocene. Ecosphere 6: 1–55.

4. Arnold PA, Noble DWA, Nicotra AB, Kearney MR, Rezende EL, Andrew SC, Briceño VF, Buckley LB, Christian KA, Clusella-Trullas S, et al. 2025. A Framework for Modelling Thermal Load Sensitivity Across Life. Global Change Biology 31: e70315.

6. Aspinwall MJ, Pfautsch S, Tjoelker MG, Vårhammar A, Possell M, Drake JE, Reich PB, Tissue DT, Atkin OK, Rymer PD, et al. 2019. Range size and growth temperature influence *Eucalyptus* species responses to an experimental heatwave. Global Change Biology 25: 1665–1684.

7. Bachofen C, D’Odorico P, Buchmann N. 2020. Light and VPD gradients drive foliar nitrogen partitioning and photosynthesis in the canopy of European beech and silver fir. Oecologia 192: 323–339.

8. Bates DM. 2018. lme4: Mixed-effects modeling with R.

9. Batllori E, Lloret F, Aakala T, Anderegg WRL, Aynekulu E, Bendixsen DP, Bentouati A, Bigler C, Burk CJ, Camarero JJ, et al. 2020. Forest and woodland replacement patterns following drought-related mortality. Proceedings of the National Academy of Sciences 117: 29720–29729.

10. Bernacchi CJ, Portis AR, Nakano H, Von Caemmerer S, Long SP. 2002. Temperature Response of Mesophyll Conductance. Implications for the Determination of Rubisco Enzyme Kinetics and for Limitations to Photosynthesis in Vivo. Plant Physiology 130: 1992–1998.

11. Berry J, Bjorkman O. 1980. Photosynthetic Response and Adaptation to Temperature in Higher Plants. Annual Review of Plant Physiology 31: 491–543.

12. Bose A, Tiwari B, Chattopadhyay MK, Gupta S, Ghosh B. 1999. Thermal stress induces differential degradation of Rubisco in heat-sensitive and heat-tolerant rice. Physiologia Plantarum 105: 89–94.

13. Breshears DD, Fontaine JB, Ruthrof KX, Field JP, Feng X, Burger JR, Law DJ, Kala J, Hardy GEStJ. 2021. Underappreciated plant vulnerabilities to heat waves. New Phytologist 231: 32–39.

14. Brodribb TJ, Powers J, Cochard H, Choat B. 2020. Hanging by a thread? Forests and drought. Science 368: 261– 266.

15. Bueno A, Alfarhan A, Arand K, Burghardt M, Deininger A-C, Hedrich R, Leide J, Seufert P, Staiger S, Riederer M. 2019. Effects of temperature on the cuticular transpiration barrier of two desert plants with water-spender and water-saver strategies. Journal of Experimental Botany 70: 1613–1625.

16. Cabrera HM, Rada F, Cavieres L. 1998. Effects of temperature on photosynthesis of two morphologically contrasting plant species along an altitudinal gradient in the tropical high Andes. Oecologia 114: 145–152.

17. Chen A, Huang L, Liu Q, Piao S. 2021. Optimal temperature of vegetation productivity and its linkage with climate and elevation on the Tibetan Plateau. Global Change Biology 27: 1942–1951.

18. Chen J, Yue K, Shen L, Zheng C, Zhu Y, Han K, Kai L. 2023. Aquaporins and CO2 diffusion across biological membrane. Frontiers in Physiology 14: 1205290.

19. Ciais Ph, Reichstein M, Viovy N, Granier A, Ogée J, Allard V, Aubinet M, Buchmann N, Bernhofer Chr, Carrara A, et al. 2005. Europe-wide reduction in primary productivity caused by the heat and drought in 2003. Nature 437: 529–533.

20. Cochard H, Martin R, Gross P. 2000. Temperature effects on hydraulic conductance and water relations of Quercus robur L. Journal of Experimental Botany 51: 1255–1259.

21. Cook AM, Rezende EL, Petrou K, Leigh A. 2024. Beyond a single temperature threshold: Applying a cumulative thermal stress framework to plant heat tolerance. Ecology Letters 27: e14416.

22. Crous KY, Uddling J, De Kauwe MG. 2022. Temperature responses of photosynthesis and respiration in evergreen trees from boreal to tropical latitudes. New Phytologist 234: 353–374.

23. Curtis EM, Knight CA, Petrou K, Leigh A. 2014. A comparative analysis of photosynthetic recovery from thermal stress: a desert plant case study. Oecologia 175: 1051–1061.

24. Deluigi J, Bachofen C, Didion-Gency M, Gisler J, Mas E, Mekarni L, Poretti A, Schaub M, Vitasse Y, Grossiord C. 2025. Prolonged warming and drought reduce canopy-level net carbon uptake in beech and oak saplings despite photosynthetic and respiratory acclimation. New Phytologist 246: 2015–2028.

25. Deutsch CA, Tewksbury JJ, Huey RB, Sheldon KS, Ghalambor CK, Haak DC, Martin PR. 2008. Impacts of climate warming on terrestrial ectotherms across latitude. Proceedings of the National Academy of Sciences 105: 6668–6672.

26. Diao H, Cernusak LA, Saurer M, Gessler A, Siegwolf RTW, Lehmann MM. 2024. Uncoupling of stomatal conductance and photosynthesis at high temperatures: mechanistic insights from online stable isotope techniques. New Phytologist 241: 2366–2378.

27. Didion-Gency M, Gauthey A, Johnson KM, Schuler P, Grossiord C. 2025. Leaf Excision and Exposure Duration Alter the Estimates of the Irreversible Photosynthetic Thermal Thresholds. Plant, Cell & Environment: 1–12.

28. Didion-Gency M, Gauthey A, Johnson KM, Schuler P, Grossiord C. 2025. Leaf Excision and Exposure Duration Alter the Estimates of the Irreversible Photosynthetic Thermal Thresholds. Plant, Cell & Environment 48: 5357– 5368.

29. Didion-Gency M, Gessler A, Buchmann N, Gisler J, Schaub M, Grossiord C. 2022. Impact of warmer and drier conditions on tree photosynthetic properties and the role of species interactions. New Phytologist 236: 547–560.

30. Doughty CE, Goulden ML. 2008. Are tropical forests near a high temperature threshold? Journal of Geophysical Research: Biogeosciences 113: 2007JG000632.

31. Downton WJS, Berry JA, Seemann JR. 1984. Tolerance of Photosynthesis to High Temperature in Desert Plants. Plant Physiology 74: 786–790.

32. Drake JE, Tjoelker MG, Vårhammar A, Medlyn BE, Reich PB, Leigh A, Pfautsch S, Blackman CJ, López R, Aspinwall MJ, et al. 2018a. Trees tolerate an extreme heatwave via sustained transpirational cooling and increased leaf thermal tolerance. Global Change Biology 24: 2390–2402.

33. Drake JE, Tjoelker MG, Vårhammar A, Medlyn BE, Reich PB, Leigh A, Pfautsch S, Blackman CJ, López R, Aspinwall MJ, et al. 2018b. Trees tolerate an extreme heatwave via sustained transpirational cooling and increased leaf thermal tolerance. Global Change Biology 24: 2390–2402.

34. Duursma RA, Blackman CJ, Lopéz R, Martin-StPaul NK, Cochard H, Medlyn BE. 2019. On the minimum leaf conductance: its role in models of plant water use, and ecological and environmental controls. New Phytologist 221: 693–705.

35. Eaton-Rye JJ, Tripathy BC, Sharkey TD (Eds). 2012. Photosynthesis: Plastid Biology, Energy Conversion and Carbon Assimilation. Dordrecht: Springer Netherlands.

36. Faber AH, Ørsted M, Ehlers BK. 2024. Application of the thermal death time model in predicting thermal damage accumulation in plants. bioRxiv.

37. Feeley K, Martinez-Villa J, Perez T, Silva Duque A, Triviño Gonzalez D, Duque A. 2020. The Thermal Tolerances, Distributions, and Performances of Tropical Montane Tree Species. Frontiers in Forests and Global Change 3: 25.

38. Fernandes VDAB, Farnese FS, Arantes BR, Fontineles Da Silva ML, Silva FG, Torres-Ruiz JM, Slot M, Cochard H, Menezes-Silva PE. 2025. Leaf minimum conductance dynamics during and after heat stress: Implications for plant survival under hotter droughts. Plant Physiology 197: kiaf026.

39. Fischer RA, Turner NC. 1978. Plant Productivity in the Arid and Semiarid Zones. Annual Review of Plant Physiology 29: 277–317.

40. Fox J, Friendly M, Weisberg S. 2013. Hypothesis Tests for Multivariate Linear Models Using the car Package. The R Journal 5: 39.

41. Fredeen AL, Sage RF. 1999. Temperature and humidity effects on branchlet gas-exchange in white spruce: an explanation for the increase in transpiration with branchlet temperature. Trees 14: 0161.

42. Garen JC, Michaletz ST. 2025. Temperature governs the relative contributions of cuticle and stomata to leaf minimum conductance. New Phytologist 245: 1911–1923.

43. Gauthey A, Bachofen C, Deluigi J, Didion-Gency M, D’Odorico P, Gisler J, Mas E, Schaub M, Schuler P, Still CJ, et al. 2023. Absence of canopy temperature variation despite stomatal adjustment in *Pinus sylvestris* under multidecadal soil moisture manipulation. New Phytologist 240: 127–137.

44. Gauthey A, Kahmen A, Limousin J, Vilagrosa A, Didion-Gency M, Mas E, Milano A, Tunas A, Grossiord C. 2024. High heat tolerance, evaporative cooling, and stomatal decoupling regulate canopy temperature and their safety margins in three European oak species. Global Change Biology 30: e17439.

45. Gayen AK. 1949. The Distribution of ‘Student’s t in Random Samples of any Size Drawn from Non-Normal Universes. Biometrika 36: 353.

46. Gazol A, Camarero JJ. 2022. Compound climate events increase tree drought mortality across European forests. Science of The Total Environment 816: 151604.

47. Giorgi F. 2006. Climate change hot-spots. Geophysical Research Letters 33: 2006GL025734.

48. Giorgi F, Bi X, Pal J. 2004. Mean, interannual variability and trends in a regional climate change experiment over Europe. II: climate change scenarios (2071-2100). Climate Dynamics 23: 839–858.

49. Giorio P. 2011. Black leaf-clips increased minimum fluorescence emission in clipped leaves exposed to high solar radiation during dark adaptation. Photosynthetica 49.

50. Gong X-W, Hao G-Y. 2023. The synergistic effect of hydraulic and thermal impairments accounts for the severe crown damage in Fraxinus mandshurica seedlings following the combined drought-heatwave stress. Science of The Total Environment 856: 159017.

51. Grossiord C, Buckley TN, Cernusak LA, Novick KA, Poulter B, Siegwolf RTW, Sperry JS, McDowell NG. 2020. Plant responses to rising vapor pressure deficit. New Phytologist 226: 1550–1566.

52. Hara C, Inoue S, Ishii HR, Okabe M, Nakagaki M, Kobayashi H. 2021. Tolerance and acclimation of photosynthesis of nine urban tree species to warmer growing conditions. Trees 35: 1793–1806.

53. Hargreaves GH, Samani ZA. 1985. Reference Crop Evapotranspiration from Temperature. Applied Engineering in Agriculture 1: 96–99.

54. Harley PC, Baldocchi DD. 1995. Scaling carbon dioxide and water vapour exchange from leaf to canopy in a deciduous forest. I. Leaf model parametrization. Plant, Cell & Environment 18: 1146–1156.

55. Holland SM. 2019. Principal components analysis (PCA).

56. Hüve K, Bichele I, Kaldmäe H, Rasulov B, Valladares F, Niinemets Ü. 2019. Responses of Aspen Leaves to Heatflecks: Both Damaging and Non-Damaging Rapid Temperature Excursions Reduce Photosynthesis. Plants 8: 145.

57. IPCC. 2023. Climate Change 2021 – The Physical Science Basis: Working Group I Contribution to the Sixth Assessment Report of the Intergovernmental Panel on Climate Change. Cambridge University Press.

58. June T, Evans JR, Farquhar GD. 2004. A simple new equation for the reversible temperature dependence of photosynthetic electron transport: a study on soybean leaf. Functional Plant Biology 31: 275.

59. Knight CA, Ackerly DD. 2002. An ecological and evolutionary analysis of photosynthetic thermotolerance using the temperature-dependent increase in fluorescence. Oecologia 130: 505–514.

60. Krause GH, Winter K, Krause B, Jahns P, García M, Aranda J, Virgo A. 2010. High-temperature tolerance of a tropical tree, Ficus insipida: methodological reassessment and climate change considerations. Functional Plant Biology 37: 890.

61. Kubien DS, Sage RF. 2008. The temperature response of photosynthesis in tobacco with reduced amounts of Rubisco. *Plant*, Cell & Environment 31: 407–418.

62. Kullberg AT, Coombs L, Soria Ahuanari RD, Fortier RP, Feeley KJ. 2024. Leaf thermal safety margins decline at hotter temperatures in a natural warming ‘experiment’ in the Amazon. New Phytologist 241: 1447–1463.

63. Kumarathunge DP, Drake JE, Tjoelker MG, López R, Pfautsch S, Vårhammar A, Medlyn BE. 2020. The temperature optima for tree seedling photosynthesis and growth depend on water inputs. Global Change Biology 26: 2544–2560.

64. Lancaster LT, Humphreys AM. 2020. Global variation in the thermal tolerances of plants. Proceedings of the National Academy of Sciences 117: 13580–13587.

65. Leon-Garcia IV, Lasso E. 2019. High heat tolerance in plants from the Andean highlands: Implications for paramos in a warmer world (J Dai, Ed.). PLOS ONE 14: e0224218.

66. Leuschner C, Ellenberg H. 2017. Ecology of Central European Forests: Vegetation Ecology of Central Europe, Volume I.Cham: Springer International Publishing.

67. Lin Y-S, Medlyn BE, Ellsworth DS. 2012. Temperature responses of leaf net photosynthesis: the role of component processes. Tree Physiology 32: 219–231.

68. Marchin RM, Broadhead AA, Bostic LE, Dunn RR, Hoffmann WA. 2016. Stomatal acclimation to vapour pressure deficit doubles transpiration of small tree seedlings with warming. *Plant*, Cell & Environment 39: 2221–2234.

69. Marchin RM, Medlyn BE, Tjoelker MG, Ellsworth DS. 2023. Decoupling between stomatal conductance and photosynthesis occurs under extreme heat in broadleaf tree species regardless of water access. Global Change Biology 29: 6319–6335.

70. Marias DE, Meinzer FC, Still C. 2017. Leaf age and methodology impact assessments of thermotolerance of Coffea arabica. Trees 31: 1091–1099.

71. Marias DE, Meinzer FC, Woodruff DR, McCulloh KA. 2016. Thermotolerance and heat stress responses of Douglas-fir and ponderosa pine seedling populations from contrasting climates (D Tissue, Ed.). Tree Physiology: treephys;tpw117v1.

72. Maxwell K, Johnson GN. 2000a. Chlorophyll fluorescence—a practical guide. Journal of Experimental Botany 51: 659–668.

73. Maxwell K, Johnson GN. 2000b. Chlorophyll fluorescence—a practical guide. Journal of Experimental Botany 51: 659–668.

74. Miller BD, Carter KR, Reed SC, Wood TE, Cavaleri MA. 2021. Only sun-lit leaves of the uppermost canopy exceed both air temperature and photosynthetic thermal optima in a wet tropical forest. Agricultural and Forest Meteorology 301–302: 108347.

75. Missik JEC, Oishi AC, Benson MC, Meretsky VJ, Phillips RP, Novick KA. 2021. Performing gas-exchange measurements on excised branches - evaluation and recommendations. Photosynthetica 59: 61–73.

76. Miyazawa Y, Tateishi M, Komatsu H, Kumagai T, Otsuki K. 2011. Are measurements from excised leaves suitable for modeling diurnal patterns of gas exchange of intact leaves? Hydrological Processes 25: 2924–2930.

77. Neuner G, Buchner O. 2023. The dose makes the poison: The longer the heat lasts, the lower the temperature for functional impairment and damage. Environmental and Experimental Botany 212: 105395.

78. Okubo N, Inoue S, Ishii H. 2023a. Tolerance and Acclimation of the Leaves of Nine Urban Tree Species to High Temperatures. Forests 14: 1639.

79. Okubo N, Inoue S, Ishii H. 2023b. Tolerance and Acclimation of the Leaves of Nine Urban Tree Species to High Temperatures. Forests 14: 1639.

80. O’sullivan OS, Heskel MA, Reich PB, Tjoelker MG, Weerasinghe LK, Penillard A, Zhu L, Egerton JJG, Bloomfield KJ, Creek D, et al. 2017. Thermal limits of leaf metabolism across biomes. Global Change Biology 23: 209–223.

81. Park Williams A, Allen CD, Macalady AK, Griffin D, Woodhouse CA, Meko DM, Swetnam TW, Rauscher SA, Seager R, Grissino-Mayer HD, et al. 2013. Temperature as a potent driver of regional forest drought stress and tree mortality. Nature Climate Change 3: 292–297.

82. Peñuelas J, Sardans J. 2021. Global Change and Forest Disturbances in the Mediterranean Basin: Breakthroughs, Knowledge Gaps, and Recommendations. Forests 12: 603.

83. Peñuelas J, Sardans J, Filella I, Estiarte M, Llusià J, Ogaya R, Carnicer J, Bartrons M, Rivas-Ubach A, Grau O, et al. 2017. Impacts of Global Change on Mediterranean Forests and Their Services. Forests 8: 463.

84. Perez TM, Socha A, Tserej O, Feeley KJ. 2021. Photosystem II heat tolerances characterize thermal generalists and the upper limit of carbon assimilation. Plant, Cell & Environment 44: 2321–2330.

85. Rao MP, Davi NK, Magney TS, Andreu-Hayles L, Nachin B, Suran B, Varuolo-Clarke AM, Cook BI, D’Arrigo RD, Pederson N, et al. 2023. Approaching a thermal tipping point in the Eurasian boreal forest at its southern margin. Communications Earth & Environment 4: 247.

86. Rey-Sánchez A, Slot M, Posada J, Kitajima K. 2016. Spatial and seasonal variation in leaf temperature within the canopy of a tropical forest. Climate Research 71: 75–89.

87. Rödenbeck C, Zaehle S, Keeling R, Heimann M. 2020. The European carbon cycle response to heat and drought as seen from atmospheric CO _2_ data for 1999–2018. Philosophical Transactions of the Royal Society B: Biological Sciences 375: 20190506.

88. Sage RF, Kubien DS. 2007. The temperature response of C_3_ and C_4_ photosynthesis. *Plant*, Cell & Environment 30: 1086–1106.

89. Salisbury FB, Spomer GG. 1964. Leaf temperatures of alpine plants in the field. Planta 60: 497–505.

90. Salvucci ME, Crafts-Brandner SJ. 2004a. Mechanism for deactivation of Rubisco under moderate heat stress. Physiologia Plantarum 122: 513–519.

91. Salvucci ME, Crafts-Brandner SJ. 2004b. Mechanism for deactivation of Rubisco under moderate heat stress. Physiologia Plantarum 122: 513–519.

92. Santiago LS, Mulkey SS. 2003. A Test of Gas Exchange Measurements on Excised Canopy Branches of Ten Tropical Tree Species. Photosynthetica 41: 343–347.

93. Sastry A, Barua D. 2017. Leaf thermotolerance in tropical trees from a seasonally dry climate varies along the slow-fast resource acquisition spectrum. Scientific Reports 7: 11246.

94. Saxton KE, Rawls WJ, Romberger JS, Papendick RI. 1986. Estimating Generalized Soil-water Characteristics from Texture. Soil Science Society of America Journal 50: 1031–1036.

95. Schönbeck LC, Schuler P, Lehmann MM, Mas E, Mekarni L, Pivovaroff AL, Turberg P, Grossiord C. 2022. Increasing temperature and vapour pressure deficit lead to hydraulic damages in the absence of soil drought. Plant, Cell & Environment 45: 3275–3289.

96. Schreiber U, Berry JA. 1977. Heat-induced changes of chlorophyll fluorescence in intact leaves correlated with damage of the photosynthetic apparatus. Planta 136: 233–238.

97. Schuster A-C, Burghardt M, Alfarhan A, Bueno A, Hedrich R, Leide J, Thomas J, Riederer M. 2016. Effectiveness of cuticular transpiration barriers in a desert plant at controlling water loss at high temperatures. AoB PLANTS 8: plw027.

98. Schymanski SJ, Or D, Zwieniecki M. 2013. Stomatal Control and Leaf Thermal and Hydraulic Capacitances under Rapid Environmental Fluctuations (W Bauerle, Ed.). PLoS ONE 8: e54231.

99. Sendall KM, Reich PB, Zhao C, Jihua H, Wei X, Stefanski A, Rice K, Rich RL, Montgomery RA. 2015. Acclimation of photosynthetic temperature optima of temperate and boreal tree species in response to experimental forest warming. Global Change Biology 21: 1342–1357.

100. Shapiro SS, Wilk MB. 1965. An Analysis of Variance Test for Normality (Complete Samples). Biometrika 52: 591.

101. Sirvydas A, Kučinskas V, Kerpauskas P, Nadzeikienė J, Kusta A. 2010. Solar radiation energy pulsations in plant leaf. Journal of environmental engineering and landscape management 18: 188–195.

102. Slot M, Cala D, Aranda J, Virgo A, Michaletz ST, Winter K. 2021. Leaf heat tolerance of 147 tropical forest species varies with elevation and leaf functional traits, but not with phylogeny. Plant, Cell & Environment 44: 2414– 2427.

103. Slot M, Krause GH, Krause B, Hernández GG, Winter K. 2019. Photosynthetic heat tolerance of shade and sun leaves of three tropical tree species. Photosynthesis Research 141: 119–130.

104. Slot M, Winter K. 2017. *In situ* temperature response of photosynthesis of 42 tree and liana species in the canopy of two Panamanian lowland tropical forests with contrasting rainfall regimes. New Phytologist 214: 1103–1117.

105. Sperlich D, Chang CT, Peñuelas J, Sabaté S. 2019. Responses of photosynthesis and component processes to drought and temperature stress: are Mediterranean trees fit for climate change? (D Way, Ed.). Tree Physiology 39: 1783–1805.

106. Tan Z-H, Zeng J, Zhang Y-J, Slot M, Gamo M, Hirano T, Kosugi Y, Da Rocha HR, Saleska SR, Goulden ML, et al. 2017. Optimum air temperature for tropical forest photosynthesis: mechanisms involved and implications for climate warming. Environmental Research Letters 12: 054022.

107. Tarvainen L, Wittemann M, Mujawamariya M, Manishimwe A, Zibera E, Ntirugulirwa B, Ract C, Manzi OJL, Andersson MX, Spetea C, et al. 2022. Handling the heat – photosynthetic thermal stress in tropical trees. New Phytologist 233: 236–250.

108. Teskey R, Wertin T, Bauweraerts I, Ameye M, Mcguire MA, Steppe K. 2015. Responses of tree species to heat waves and extreme heat events. Plant, Cell & Environment 38: 1699–1712.

109. Thebud R, Santarius KA. 1982. Effects of High-Temperature Stress on Various Biomembranes of Leaf Cells *In Situ* and *In Vitro*. Plant Physiology 70: 200–205.

110. Uehlein N, Otto B, Hanson DT, Fischer M, McDowell N, Kaldenhoff R. 2008. Function of *Nicotiana tabacum* Aquaporins as Chloroplast Gas Pores Challenges the Concept of Membrane CO_2_ Permeability. The Plant Cell 20: 648–657.

111. Urban J, Ingwers M, McGuire MA, Teskey RO. 2017. Stomatal conductance increases with rising temperature. Plant Signaling & Behavior 12: e1356534.

112. Vogel S. 1970. Convective Cooling at Low Airspeeds and the Shapes of Broad Leaves. Journal of Experimental Botany 21: 91–101.

113. Von Caemmerer S, Evans JR. 2015. Temperature responses of mesophyll conductance differ greatly between species. Plant, Cell & Environment 38: 629–637.

114. Wahid A, Gelani S, Ashraf M, Foolad M. 2007. Heat tolerance in plants: An overview. Environmental and Experimental Botany 61: 199–223.

115. Wise RR, Olson AJ, Schrader SM, Sharkey TD. 2004. Electron transport is the functional limitation of photosynthesis in field-grown Pima cotton plants at high temperature. Plant, Cell & Environment 27: 717–724.

116. Yamori W, Hikosaka K, Way DA. 2014. Temperature response of photosynthesis in C3, C4, and CAM plants: temperature acclimation and temperature adaptation. Photosynthesis Research 119: 101–117.

117. Yordanov I, Dilova S, Petkova R, Pangelova T, Goltsev V, Suss K-H. 1986. Mechanisms of the temperature damage and acclimation of the photosynthetic apparatus. Photobiochemistry and Photobiophysics 12: 147–155.

118. Zhou Y, Kitudom N, Fauset S, Slot M, Fan Z, Wang J, Liu W, Lin H. 2023. Leaf thermal regulation strategies of canopy species across four vegetation types along a temperature and precipitation gradient. Agricultural and Forest Meteorology 343: 109766.

119. Zhu L, Bloomfield KJ, Hocart CH, Egerton JJG, O’Sullivan OS, Penillard A, Weerasinghe LK, Atkin OK. 2018. Plasticity of photosynthetic heat tolerance in plants adapted to thermally contrasting biomes. Plant, Cell & Environment 41: 1251–1262.

120. Zhu Y, Kang H, Xie Q, Wang Z, Yin S, Liu C. 2012. Pattern of leaf vein density and climate relationship of Quercus variabilis populations remains unchanged with environmental changes. Trees 26: 597–607.

121. Zhu L, Li H, Thorpe MR, Hocart CH, Song X. 2021. Stomatal and mesophyll conductance are dominant limitations to photosynthesis in response to heat stress during severe drought in a temperate and a tropical tree species. Trees 35: 1613–1626.

