## Supplementary Information for "Living on the edge: warmer climate reduces leaf thermal safety margins and causes gas exchange decoupling in Mediterranean shrubs"

**ORCIDs**:

Margaux Didion-Gency: https://orcid.org/0000-0001-89673655

Pol Soler: https://orcid.org/0009-0001-7072-1992

Eva Castells: https://orcid.org/0000-0001-7423-2742

Jordi Martinez-Vilalta: https://orcid.org/0000-0002-2332-7298

19 **Table S1**: Shrubs information across species and study sites.

| Location | Site code | Species | Replicates | Mean H (cm) | Mean BD (mm) | Mean $N_{stem}$ | Mean $P_{competition}$ (%) | Mean $P_{crown}$ (%) |
|---|---|---|---|---|---|---|---|---|
| Pas de l'Ase | PAS | Au | 8 | 211.00 | 34.46 | 6.88 | 65.00 | 3.67 |
| | | Pl | 8 | 168.69 | 47.62 | 2.00 | 40.00 | 2.32 |
| | | Ra | 7 | 219.86 | 26.35 | 2.57 | 46.43 | 0.90 |
| | | Sr | 8 | 140.75 | 32.19 | 5.88 | 33.13 | 1.29 |
| Alòs de Balaguer | ALO | Ao | 8 | 196.13 | 13.96 | 16.25 | 61.88 | 2.23 |
| | | Bs | 8 | 146.75 | 18.52 | 12.50 | 38.75 | 0.88 |
| | | Ra | 8 | 164.00 | 20.92 | 2.13 | 70.00 | 0.59 |
| | | Sr | 8 | 95.13 | 19.57 | 6.38 | 46.25 | 1.05 |
| Garraf | GAR | Au | 8 | 212.50 | 40.20 | 16.38 | 54.38 | 4.04 |
| | | Bs | 8 | 163.38 | 28.32 | 5.63 | 53.13 | 1.00 |
| | | Pl | 7 | 127.43 | 25.36 | 9.86 | 73.57 | 1.59 |
| | | Ra | 8 | 168.25 | 32.01 | 3.75 | 62.50 | 1.21 |
| | | Sr | 7 | 98.88 | 20.60 | 8.88 | 34.38 | 0.83 |
| Cingles de Bertí | CIN | Au | 9 | 275.00 | 39.68 | 6.78 | 81.67 | 5.60 |
| | | Pl | 8 | 119.38 | 17.48 | 16.13 | 68.75 | 4.28 |
| | | Sr | 8 | 118.75 | 18.33 | 5.75 | 43.75 | 0.95 |
| Montsec | MON | Ao | 7 | 308.29 | 20.07 | 23.71 | 72.86 | 5.00 |
| | | Bs | 7 | 133.14 | 21.79 | 19.29 | 24.29 | 1.68 |
| | | Ra | 8 | 123.50 | 25.33 | 1.38 | 75.00 | 0.40 |
| | | Sr | 8 | 94.00 | 15.95 | 19.50 | 45.00 | 1.76 |
| Pentina | PEN | Ao | 9 | 366.89 | 42.73 | 15.67 | 43.33 | 5.64 |
| | | Bs | 8 | 185.75 | 21.15 | 8.75 | 23.75 | 1.49 |

20 Abbreviation: Ao: *A. ovalis*, Au: *A. unedo*, Bs: *B. sempervirens*, Pl: *P. lentiscus*, Ra: *R. alaternus*, and Sr: *S. rosmarinus*.

**Table S2**: Environmental conditions during measurement campaigns across study sites.

| Location | Site code | Mean $T_{air}$ (°C) | $\Delta T_{air,max}$ (°C) | Δdays to $T_{air,max}$ (days) | Mean RH (%) | Mean VPD (kPa) | Mean PPFD (µmol $m^{-2}$ $s^{-1}$) | Mean wind speed (m $s^{-1}$) |
|---|---|---|---|---|---|---|---|---|
| Pas de l'Ase | PAS | 28.35 (19.2 – 34.8) | 4.1 | -16 | 58.35 (40.8 – 98.7) | 1.61 | 1153 | 0.78 |
| Alòs de Balaguer | ALO | 27.08 (17.2 – 34.3) | 2.6 | -12 | 55.59 (40.3 – 94.8) | 1.59 | 1222 | 1.23 |
| Garraf | GAR | 26.53 (18.0 – 32.3) | 3.7 | -7 | 61.37 (40.7 – 100) | 1.34 | 1246 | 1.06 |
| Cingles de Bertí | CIN | 28.64 (17.7 – 35.6) | 0.1 | -2 | 42.03 (27.2 – 83.5) | 2.27 | 1368 | 0.63 |
| Montsec | MON | 21.76 (13.3 – 27.7) | 4.0 | 3 | 79.69 (57.0 – 100) | 0.53 | 1166 | 1.12 |
| Pentina | PEN | 22.50 (15.6 – 27.8) | 2.6 | 8 | 82.80 (60.0 – 100) | 0.47 | 727 | 0.77 |

[1]Abbreviation: $T_{air}$: air temperature, $\Delta T_{air,max}$: difference between measurement-day $T_{air,ma}$x and yearly $T_{air,max}$; Δdays to $T_{air,max}$: number of days between measurement-day and yearly $T_{air,max}$; RH : relative humidity, VPD: vapor pressure deficit, PPFD: photosynthetic photon flux density.

[2]Daily meteorological conditions were obtained from Meteo.cat stations where site-specific downscaling was performed.

**Table S3**: Soil characteristics of the study sites.

| Location | Site code | Organic carbon (%) | Organic matter (%) | Organic nitrogen (%) | Nitrate (%) | Phosphorus (%) | Clay (%) | Silt (%) | Sand (%) | *b* | x-axis from PCA |
|---|---|---|---|---|---|---|---|---|---|---|---|
| Pas de l'Ase | PAS | 1.22 | 2.1 | 0.10 | 2.5 | < 5 | 21.3 | 37.8 | 41.0 | -5.39 | 3.48 |
| Alòs de Balaguer | ALO | 2.26 | 3.9 | 0.25 | 2.1 | < 5 | 15.4 | 29.0 | 55.6 | -5.33 | 1.16 |
| Garraf | GAR | 4.41 | 7.6 | 0.53 | 2.9 | < 5 | 21.5 | 19.2 | 59.3 | -6.80 | -1.54 |
| Cingles de Bertí | CIN | 2.49 | 4.3 | 0.19 | 2.0 | < 5 | 19.4 | 25.2 | 55.4 | -6.05 | 0.86 |
| Montsec | MON | 4.81 | 8.3 | 0.52 | 2.1 | < 5 | 21.3 | 18.6 | 60.1 | -6.83 | -1.89 |
| Pentina | PEN | 5.51 | 9.5 | 0.49 | 2.0 | < 5 | 11.3 | 22.5 | 66.2 | -5.15 | -2.08 |

Abbreviation: *b*: coefficient from the Saxton equation.

**Table S4:** Principal Component Analysis (PCA) of the soil variables, including organic carbon (OC), organic matter (OM), organic nitrogen (ON), nitrate ($NO_3$), clay, silt, sand and *b* coefficient from the Saxton equation (*b*): eigenvalues, cumulated variance explained ($variance_{cum}$), variable loadings, and contributions. Variables selected for the models are highlighted in bold.

| Sources of variations | PC | Eigenvalue | $Variance_{cum}$ (%) | Loading | Contribution |
|---|---|---|---|---|---|
| OC | PC1 | 4.90 | 61.2 | -0.44 | 3.99 |
| | PC2 | 2.29 | 89.8 | -0.05 | 0.11 |
| OM | PC1 | 4.90 | 61.2 | -0.44 | 3.99 |
| | PC2 | 2.29 | 89.8 | -0.05 | 0.11 |
| ON | PC1 | 4.90 | 61.2 | -0.43 | 3.86 |
| | PC2 | 2.29 | 89.8 | 0.10 | 0.48 |
| $NO_3$ | PC1 | 4.90 | 61.2 | 0.03 | 0.02 |
| | PC2 | 2.29 | 89.8 | 0.50 | 11.06 |
| Clay | PC1 | 4.90 | 61.2 | 0.12 | 0.28 |
| | PC2 | 2.29 | 89.8 | 0.61 | 16.48 |
| Silt | PC1 | 4.90 | 61.2 | 0.43 | 3.75 |
| | PC2 | 2.29 | 89.8 | -0.13 | 0.69 |
| Sand | PC1 | 4.90 | 61.2 | -0.42 | 3.64 |
| | PC2 | 2.29 | 89.8 | -0.20 | 1.66 |
| ***b*** | **PC1** | **4.90** | **61.2** | **0.21** | **0.90** |
| | **PC2** | **2.29** | **89.8** | **-0.55** | **13.06** |

**Table S5:** Principal components analysis (PCA) of the models fixed effects, including the climatic variables (i.e., maximum monthly air temperature, MMAT 2009-2024; MMAT 2024; air maximum temperature, $T_{air,max}$; air minimum temperature, $T_{air,min}$; P/PET), the soil variables (i.e., *b* coefficient from the Saxton equation, *b*; x-axis from the soil PCA; organic carbon, OC; organic matter, OM; organic nitrogen, ON; nitrate, $NO_3$; clay; silt; sand), the individuals variables (i.e., height, H; basal diameter, BD; stem number, $N_{stem}$; plant competition, $P_{competition}$; plant crown, $P_{crown}$): eigenvalues, cumulated variance explained ($variance_{cum}$), variable loadings, and contributions. Variables selected for the models are highlighted in bold.

| Sources of variations | PC | Eigenvalue | $Variance_{cum}$ (%) | Loading | Contribution |
|---|---|---|---|---|---|
| MMAT 2009-2024 | PC1 | 10.21 | 53.7 | 0.30 | 0.89 |
| | PC2 | 2.96 | 69.3 | -0.12 | 0.47 |
| **MMAT 2024** | **PC1** | **10.21** | **53.7** | **0.30** | **0.86** |
| | **PC2** | **2.96** | **69.3** | **-0.13** | **0.58** |
| $T_{air,max}$ | PC1 | 10.21 | 53.7 | 0.30 | 0.87 |
| | PC2 | 2.96 | 69.3 | -0.12 | 0.52 |
| $T_{air,min}$ | PC1 | 10.21 | 53.7 | 0.29 | 0.84 |
| | PC2 | 2.96 | 69.3 | -0.15 | 0.74 |
| P/PET | PC1 | 10.21 | 53.7 | -0.29 | 0.83 |
| | PC2 | 2.96 | 69.3 | 0.14 | 0.69 |
| ***b*** | **PC1** | **10.21** | **53.7** | **0.15** | **0.23** |
| | **PC2** | **2.96** | **69.3** | **0.45** | **6.98** |
| x-axis | PC1 | 10.21 | 53.7 | 0.30 | 0.88 |
| | PC2 | 2.96 | 69.3 | 0.14 | 0.68 |
| OC | PC1 | 10.21 | 53.7 | -0.30 | 0.90 |
| | PC2 | 2.96 | 69.3 | -0.10 | 0.35 |
| OM | PC1 | 10.21 | 53.7 | -0.30 | 0.90 |
| | PC2 | 2.96 | 69.3 | -0.10 | 0.35 |
| ON | PC1 | 10.21 | 53.7 | -0.28 | 0.74 |
| | PC2 | 2.96 | 69.3 | -0.23 | 1.81 |
| $NO_3$ | PC1 | 10.21 | 53.7 | 0.06 | 0.04 |
| | PC2 | 2.96 | 69.3 | -0.45 | 6.95 |
| Clay | PC1 | 10.21 | 53.7 | 0.07 | 0.05 |
| | PC2 | 2.96 | 69.3 | -0.45 | 6.86 |
| Silt | PC1 | 10.21 | 53.7 | 0.28 | 0.76 |
| | PC2 | 2.96 | 69.3 | 0.21 | 1.54 |

| | | | | | |
|---|---|---|---|---|---|
| Sand | PC1 | 10.21 | 53.7 | -0.29 | 0.82 |
| | PC2 | 2.96 | 69.3 | 0.01 | 0.00 |
| **H** | **PC1** | **10.21** | **53.7** | **-0.06** | **0.03** |
| | **PC2** | **2.96** | **69.3** | **0.30** | **2.98** |
| BD | PC1 | 10.21 | 53.7 | 0.03 | 0.01 |
| | PC2 | 2.96 | 69.3 | 0.06 | 0.10 |
| $N_{stem}$ | PC1 | 10.21 | 53.7 | -0.12 | 0.13 |
| | PC2 | 2.96 | 69.3 | 0.09 | 0.26 |
| **$P_{competition}$** | **PC1** | **10.21** | **53.7** | **0.01** | **0.00** |
| | **PC2** | **2.96** | **69.3** | **-0.04** | **0.05** |
| $P_{crown}$ | PC1 | 10.21 | 53.7 | -0.05 | 0.02 |
| | PC2 | 2.96 | 69.3 | 0.24 | 1.89 |

**Table S6:** Summary of the linear effects models (slope, standard error, SE, and p-values) where the relationship of air temperature ($T_{air}$) were evaluated on the difference between leaf and air temperature ($\Delta T$). Information on whether slopes differ for each species among sites (ANOVA) is also provided. Significant effects ($p \leq 0.05$) are highlighted in bold.

| Species | Site | Location | Slope ($\Delta T \sim T_{air}$) | SE | p-value | ANOVA p-value |
|---|---|---|---|---|---|---|
| | ALO | Alòs de Balaguer | 0.07 | 0.01 | **0.000** | 0.960 |
| Ao | MON | Montsec | 0.18 | 0.01 | **0.000** | 0.960 |
| | PEN | Pentina | 0.04 | 0.01 | **0.000** | 0.960 |
| | PAS | Pas de l'Ase | 0.07 | 0.01 | **0.000** | 0.616 |
| Au | GAR | Garraf | 0.39 | 0.01 | **0.000** | 0.616 |
| | CIN | Cingles de Bertí | 0.20 | 0.01 | **0.000** | 0.616 |
| | ALO | Alòs de Balaguer | 0.08 | 0.01 | **0.000** | 0.740 |
| Bs | GAR | Garraf | 0.27 | 0.01 | **0.000** | 0.740 |
| | MON | Montsec | 0.15 | 0.01 | **0.000** | 0.740 |
| | PEN | Pentina | 0.08 | 0.01 | **0.000** | 0.740 |
| | PAS | Pas de l'Ase | 0.19 | 0.01 | **0.000** | 0.860 |
| Pl | GAR | Garraf | 0.26 | 0.01 | **0.000** | 0.860 |
| | CIN | Cingles de Bertí | 0.13 | 0.01 | **0.000** | 0.860 |
| | PAS | Pas de l'Ase | 0.08 | 0.00 | **0.000** | 0.733 |
| Ra | ALO | Alòs de Balaguer | -0.01 | 0.01 | 0.343 | 0.733 |
| | GAR | Garraf | 0.23 | 0.01 | **0.000** | 0.733 |
| | MON | Montsec | 0.10 | 0.01 | **0.000** | 0.733 |
| | PAS | Pas de l'Ase | 0.09 | 0.01 | **0.000** | 0.433 |
| | ALO | Alòs de Balaguer | 0.06 | 0.00 | **0.000** | 0.433 |
| Sr | GAR | Garraf | 0.15 | 0.01 | **0.000** | 0.433 |
| | CIN | Cingles de Bertí | 0.09 | 0.00 | **0.000** | 0.433 |
| | MON | Montsec | 0.12 | 0.01 | **0.000** | 0.433 |

Abbreviation: Ao: *A. ovalis*, Au: *A. unedo*, Bs: *B. sempervirens*, Pl: *P. lentiscus*, Ra: *R. alaternus*, and Sr: *S. rosmarinus*.

**Table S7**: Summary of the linear mixed-effects models (estimates and p-values) where the interactive effects of leaf stomatal conductance ($g_s$) and leaf temperature ($T_{leaf}$) or leaf VPD ($VPD_{leaf}$), and the additive effects of plant competition ($P_{competition}$), height (H), and *b* coefficient from the Saxton equation (*b*) were evaluated on the leaf assimilation (A). Significant effects ($p \leq 0.05$) are highlighted in bold. Information on the site and species random effect variances, the proportion of explained variance by fixed effects ($R^2$ marginal) and explained variance by fixed and random effects ($R^2$ conditional) are also provided. Model fit with the number of parameters (npar), Akaike Information Criterion (AIC), Bayesian Information Criterion (BIC), likelihood ratio tests (Chisq, Df, Pr (> Chisq)) are also shown.

| **Sources of variations** | **A** | **Sources of variations** | **A** |
|---|---|---|---|
| Fixed parts | | Fixed parts | |
| $g_s$ | **0.06 (< 0.001)** | $g_s$ | **0.08 (< 0.001)** |
| $T_{leaf}$ | **-0.04 (< 0.001)** | $VPD_{leaf}$ | **-0.15 (< 0.001)** |
| $g_s : T_{leaf}$ | **0.00 (0.002)** | $g_s : VPD_{leaf}$ | **0.01 (0.005)** |
| $P_{competition}$ | **0.00 (0.026)** | $P_{competition}$ | **-0.01 (0.021)** |
| H | **0.00 (0.003)** | H | **0.00 (0.003)** |
| *b* | 0.01 (0.978) | *b* | 0.00 (0.999) |
| Random part | | Random part | |
| Site | 1.40 | Site | 1.37 |
| Species | 0.80 | Species | 0.81 |
| $R^2$ marginal | 0.34 | $R^2$ marginal | 0.34 |
| $R^2$ conditional | 0.69 | $R^2$ conditional | 0.69 |
| Observations | 982 | Observations | 982 |

*Full linear model 1: lmer(A ~ $g_s$ + $T_{leaf}$ + $g_s : T_{leaf}$ + $P_{competition}$ + H + b + (1|Site) + (1|Species))*

*Full linear model 2: lmer(A ~ $g_s$ + $VPD_{leaf}$ + $g_s : VPD_{leaf}$ + $P_{competition}$ + H + b + (1|Site) + (1|Species))*

| | npar | AIC | BIC | Chisq | Df | Pr (> Chisq) |
|---|---|---|---|---|---|---|
| Model 1 | 10 | **3527.1** | 3576.0 | | | |
| Model 2 | 10 | 3523.6 | 3572.5 | 3.5112 | 0 | |

**Table S8:** Summary of the linear mixed-effects models (estimates and p-values) where the interactive effects of leaf transpiration (E) and leaf temperature ($T_{leaf}$) or leaf VPD ($VPD_{leaf}$), and the additive effects of plant competition ($P_{competition}$), height (H), and *b* coefficient from the Saxton equation (*b*) were evaluated on the leaf assimilation (A). Significant effects ($p \leq 0.05$) are highlighted in bold. Information on the site and species random effect variances, the proportion of explained variance by fixed effects ($R^2$ marginal) and explained variance by fixed and random effects ($R^2$ conditional) are also provided. Models fits with the number of parameters (npar), Akaike Information Criterion (AIC), Bayesian Information Criterion (BIC), likelihood ratio tests (Chisq, Df, Pr (> Chisq)) are also shown.

| **Sources of variations** | **A** | **Sources of variations** | **A** |
|---|---|---|---|
| Fixed parts | | Fixed parts | |
| E | **7.38 (< 0.001)** | E | **4.40 (< 0.001)** |
| $T_{leaf}$ | **-0.02 (0.003)** | $VPD_{leaf}$ | **-0.15 (< 0.001)** |
| E : $T_{leaf}$ | **-0.14 (< 0.001)** | E : $VPD_{leaf}$ | **-0.40 (< 0.001)** |
| $P_{competition}$ | **0.00 (0.031)** | $P_{competition}$ | **-0.01 (0.015)** |
| H | **0.00 (0.025)** | H | **0.00 (0.007)** |
| *b* | 0.07 (0.896) | *b* | 0.02 (0.971) |
| Random part | | Random part | |
| Site | 1.44 | Site | 1.39 |
| Species | 0.77 | Species | 0.79 |
| $R^2$ marginal | 0.37 | $R^2$ marginal | 0.36 |
| $R^2$ conditional | 0.72 | $R^2$ conditional | 0.71 |
| Observations | 982 | Observations | 982 |

*Full linear model 1: lmer(A ~ E +* $T_{leaf}$ *+ E:* $T_{leaf}$ *+* $P_{competition}$ *+ H + b + (1|Site) + (1|Species))*

*Full linear model 2: lmer(A ~ E +* $VPD_{leaf}$ *+ E :* $VPD_{leaf}$ *+* $P_{competition}$ *+ H + b + (1|Site) + (1|Species))*

| | npar | AIC | BIC | Chisq | Df | Pr (> Chisq) |
|---|---|---|---|---|---|---|
| Model 1 | 10 | **3446.5** | 3495.4 | | | |
| Model 2 | 10 | 3458.1 | 3507.0 | 0 | 0 | |

**Table S9**: Summary of the linear or quadratic mixed-effects models (estimates and p-values) where the additive effects of maximum monthly air temperature (MMAT) for 2024, plant competition ($P_{competition}$), height (H), and *b* coefficient from the Saxton equation (*b*) were evaluated on the photosynthetic efficiency of the photosystem II ($F_v/F_m$), critical leaf temperature ($T_{crit}$), temperature causing a 50% reduction of $F_v/F_m$ ($T_{50}$), maximum temperature ($T_{max}$), leaf optimal assimilation ($A_{opt}$), leaf optimal temperature ($T_{opt}$), and leaf thermal safety margin (TSM). Significant effects ($p \leq 0.05$) are highlighted in bold. Information on the random effect variances (site and species), the proportion of explained variance by fixed effects ($R^2$ marginal) and explained variance by fixed and random effects ($R^2$ conditional) are also provided. Models fits (Akaike Information Criterion, AIC, and p-values) are also shown.

| **Sources of variations** | **$F_v/F_m$** | **$T_{crit}$** | **Log $T_{50}$** | **Log $T_{max}$** | **Log $A_{opt}$** | **$T_{opt}$** | **TSM** |
|---|---|---|---|---|---|---|---|
| Fixed parts | | | | | | | |
| MMAT | **0.29 (0.018)** | 0.14 (0.577) | 0.17 (0.090) | **0.16 (0.005)** | -0.15 (0.174) | **0.42 (0.050)** | **-1.05 (< 0.001)** |
| I($MMAT^2$) | **-0.01 (0.019)** | - | 0.00 (0.099) | **-0.00 (0.006)** | - | - | - |
| $P_{competition}$ | 0.00 (0.413) | 0.01 (0.311) | 0.00 (0.054) | **0.00 (0.039)** | 0.00 (0.766) | -0.03 (0.082) | 0.01 (0.384) |
| H | 0.00 (0.141) | 0.00 (0.704) | 0.00 (0.338) | 0.00 (0.161) | 0.00 (0.775) | 0.00 (0.869) | 0.00 (0.465) |
| *b* | 0.03 (0.185) | -0.12 (0.897) | 0.01 (0.513) | 0.01 (0.456) | 0.09 (0.836) | 0.47 (0.548) | -0.19 (0.851) |
| Random part | | | | | | | |
| Site | 0.00 | 2.43 | 0.02 | 0.00 | 0.49 | 0.70 | 2.51 |
| Species | 0.00 | 0.81 | 0.02 | 0.00 | 0.74 | 4.41 | 0.26 |
| $R^2$ marginal | 0.11 | 0.02 | 0.18 | 0.19 | 0.07 | 0.07 | 0.56 |
| $R^2$ conditional | 0.43 | 0.53 | 0.57 | 0.42 | 0.68 | 0.26 | 0.77 |
| Observations | 173 | 173 | 173 | 173 | 172 | 172 | 173 |
| *Full linear model: lmer(traits ~ MMAT 2024 + $P_{competition}$ + H + b + (1\|Site) + (1\|Species))* | | | | | | | |
| *Full quadratic model: lmer(traits ~ MMAT 2024 + I($MMAT^2$) + $P_{competition}$ + H +b + (1\|Site) + (1\|Species))* | | | | | | | |
| Linear model | -407.3 | **718.1** | -673.2 | -629.6 | **460.6** | **1025.5** | **714.8** |
| Quadratic model | **-412.1 (0.009)** | 717.2 (0.091) | **-676.1 (0.029)** | **-636.2 (0.003)** | 458.7 (0.060) | 1024.0 (0.062) | 713.0 (0.052) |

**Table S10**: Summary of the linear mixed-effects models (estimates and p-values) where the additive effects of air temperature ($T_{air}$) for each species (i.e., Ao, Au, Bs, Pl, Ra, Sr), plant competition ($P_{competition}$), height (H), and *b* coefficient from the Saxton equation (*b*) were evaluated on the leaf temperature ($T_{leaf}$) and the difference between leaf and air temperature (ΔT). Significant effects ($p \leq 0.05$) are highlighted in bold. Information on the site random effect variances, the proportion of explained variance by fixed effects ($R^2$ marginal) and explained variance by fixed and random effects ($R^2$ conditional) are also provided.

| **Sources of variations** | $\mathbf{T_{leaf}}$ | **ΔT** |
|---|---|---|
| Fixed parts | | |
| $T_{air}$ : Ao | **1.13 (< 0.001)** | **0.13 (< 0.001)** |
| $T_{air}$ : Au | **1.14 (< 0.001)** | **0.14 (< 0.001)** |
| $T_{air}$ : Bs | **1.14 (< 0.001)** | **0.14 (< 0.001)** |
| $T_{air}$ : Pl | **1.13 (< 0.001)** | **0.13 (< 0.001)** |
| $T_{air}$ : Ra | **1.13 (< 0.001)** | **0.13 (< 0.001)** |
| $T_{air}$ : Sr | **1.13 (< 0.001)** | **0.13 (< 0.001)** |
| $P_{competition}$ | **0.00 (0.002)** | **0.00 (0.002)** |
| H | 0.00 (0.564) | 0.00 (0.564) |
| *b* | -0.37 (0.249) | -0.37 (0.249) |
| Random part | | |
| Site | 0.22 | 0.22 |
| $R^2$ marginal | 0.97 | 0.30 |
| $R^2$ conditional | 0.97 | 0.40 |
| Observations | 18431 | 18431 |

*Full linear model: lmer(traits ~ $T_{air}$ : Species + $P_{competition}$ + H + b + (1|Site))*

Abbreviation: Ao: *A. ovalis*, Au: *A. unedo*, Bs: *B. sempervirens*, Pl: *P. lentiscus*, Ra: *R. alaternus*, and Sr: *S. rosmarinus*.

**Table S11**: Summary of the linear mixed-effects models (estimates and p-values) where the interactive effects of leaf stomatal conductance ($g_s$) or leaf transpiration (E) and leaf temperature ($T_{leaf}$), and the additive effects of leaf VPD residuals ($VPD_{leaf,resid}$), plant competition ($P_{competition}$), height (H), and *b* coefficient from the Saxton equation (*b*) were evaluated on the leaf assimilation (A). Significant effects ($p \leq 0.05$) are highlighted in bold. Information on the site and species random effect variances, the proportion of explained variance by fixed effects ($R^2$ marginal) and explained variance by fixed and random effects ($R^2$ conditional) are also provided.

| **Sources of variations** | **A** | **Sources of variations** | **A** |
|---|---|---|---|
| Fixed parts | | Fixed parts | |
| $g_s$ | **0.05 (< 0.001)** | E | **7.03 (< 0.001)** |
| $T_{leaf}$ | **-0.04 (< 0.001)** | $T_{leaf}$ | **-0.03 (0.001)** |
| $g_s : T_{leaf}$ | **-0.00 (0.001)** | $E : T_{leaf}$ | **-0.13 (< 0.001)** |
| $VPD_{leaf,resid}$ | **-0.33 (< 0.001)** | $VPD_{leaf,resid}$ | **-0.24 (0.011)** |
| $P_{competition}$ | **-0.01 (0.014)** | $P_{competition}$ | **0.00 (0.020)** |
| H | **0.00 (0.002)** | H | **0.00 (0.018)** |
| *b* | -0.03 (0.949) | *b* | 0.03 (0.949) |
| Random part | | Random part | |
| Site | 1.39 | Site | 1.39 |
| Species | 0.82 | Species | 0.79 |
| $R^2$ marginal | 0.34 | $R^2$ marginal | 0.37 |
| $R^2$ conditional | 0.69 | $R^2$ conditional | 0.71 |
| Observations | 982 | Observations | 982 |

*Full linear model: lmer(A ~ $g_s + T_{leaf} + g_s : T_{leaf} + VPD_{leaf,resid} + P_{competition} + H + b$ + (1|Site) + (1|Species))*

*Full linear model: lmer(A ~ $E + T_{leaf} + E : T_{leaf} + VPD_{leaf,resid} + P_{competition} + H + b$ + (1|Site) + (1|Species))*

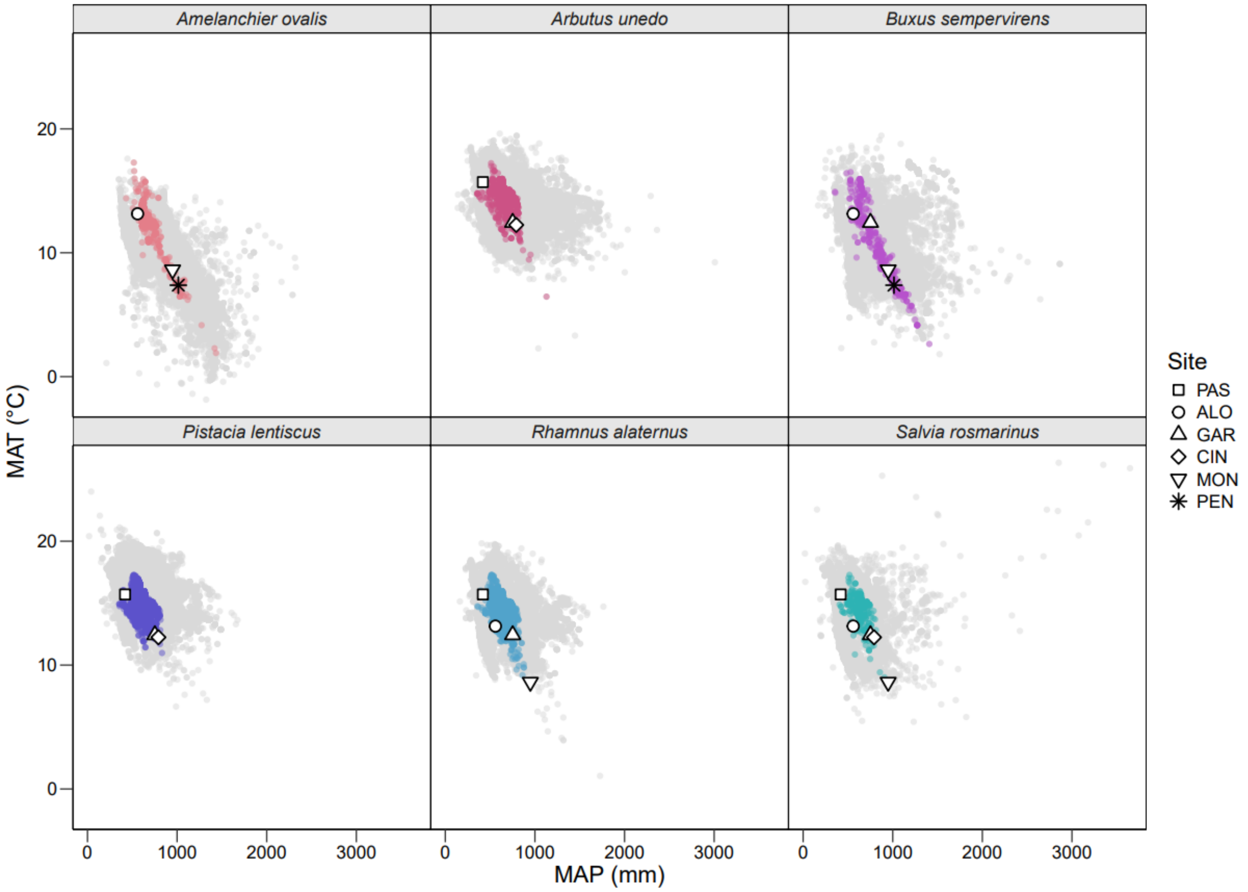


**Figure S1**: Ecological niche diagram for each species (i.e., *A. ovalis*, *A. unedo*, *B. sempervirens*, *P. lentiscus*, *R. alaternus*, and *S. rosmarinus*) comparing the mean annual precipitation (MAP) and mean annual temperature (MAT). The global distribution range of the species is highlighted in grey. The observed presence of the species in Catalonia (NE, Spain) is highlighted by color (i.e., light pink, pink, magenta, purple, blue, and cyan, respectively), and by shape for each site (i.e., PAS: Pas de l'Ase, ALO: Alòs de Balaguer, GAR: Garraf, CIN: Cingles de Bertí, MON: Montsec, PEN: Pentina).

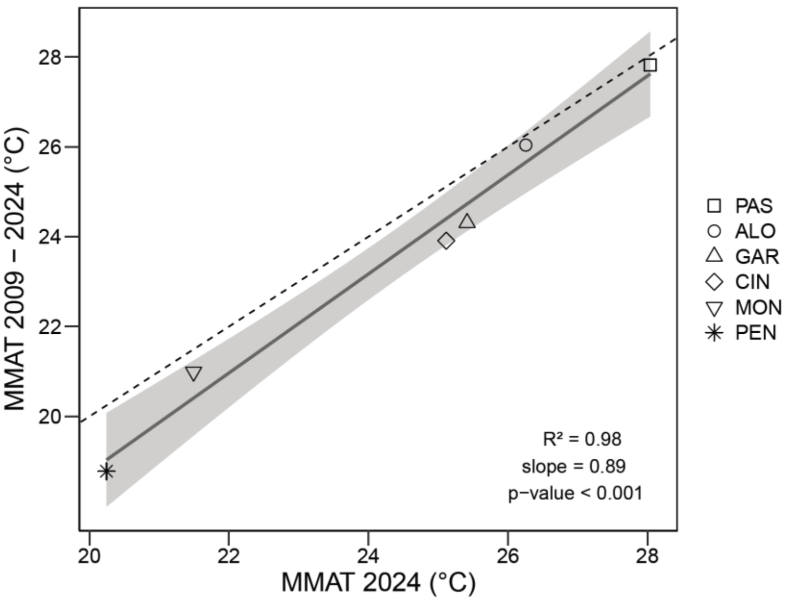


**Figure S2**: Long *vs*. short term maximum monthly air temperature (MMAT) across site (i.e., PAS: Pas de l'Ase, ALO: Alòs de Balaguer, GAR: Garraf, CIN: Cingles de Bertí, MON: Montsec, PEN: Pentina). The dashed line represents the 1:1 relationship, while the solid grey line represents the linear relationship between MMAT for 2009-2024 and MMAT for 2024. The $R^2$, slope and p-value indicate the significance of the relationship.

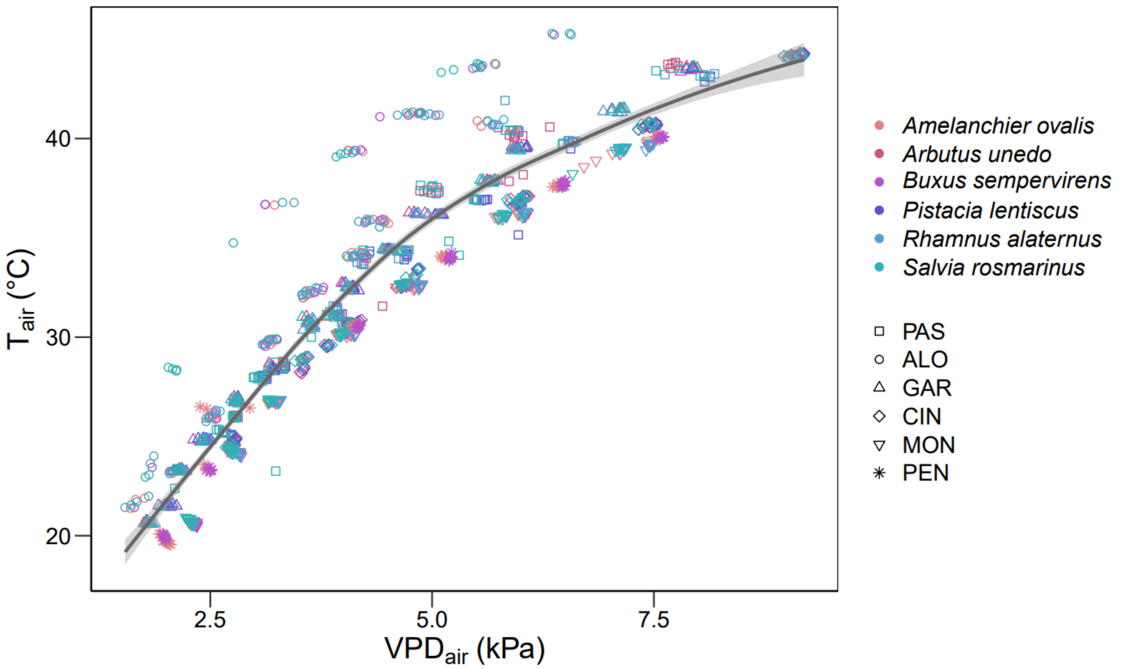


**Figure S3**: Relationship between air temperature ($T_{air}$) and air vapor pressure deficit ($VPD_{air}$) inside the LI-COR chamber for each species (i.e., *A. ovalis*, *A. unedo*, *B. sempervirens*, *P. lentiscus*, *R. alaternus*, and *S. rosmarinus*) and site (i.e., PAS: Pas de l'Ase, ALO: Alòs de Balaguer, GAR: Garraf, CIN: Cingles de Bertí, MON: Montsec, PEN: Pentina). Loess regression is shown.

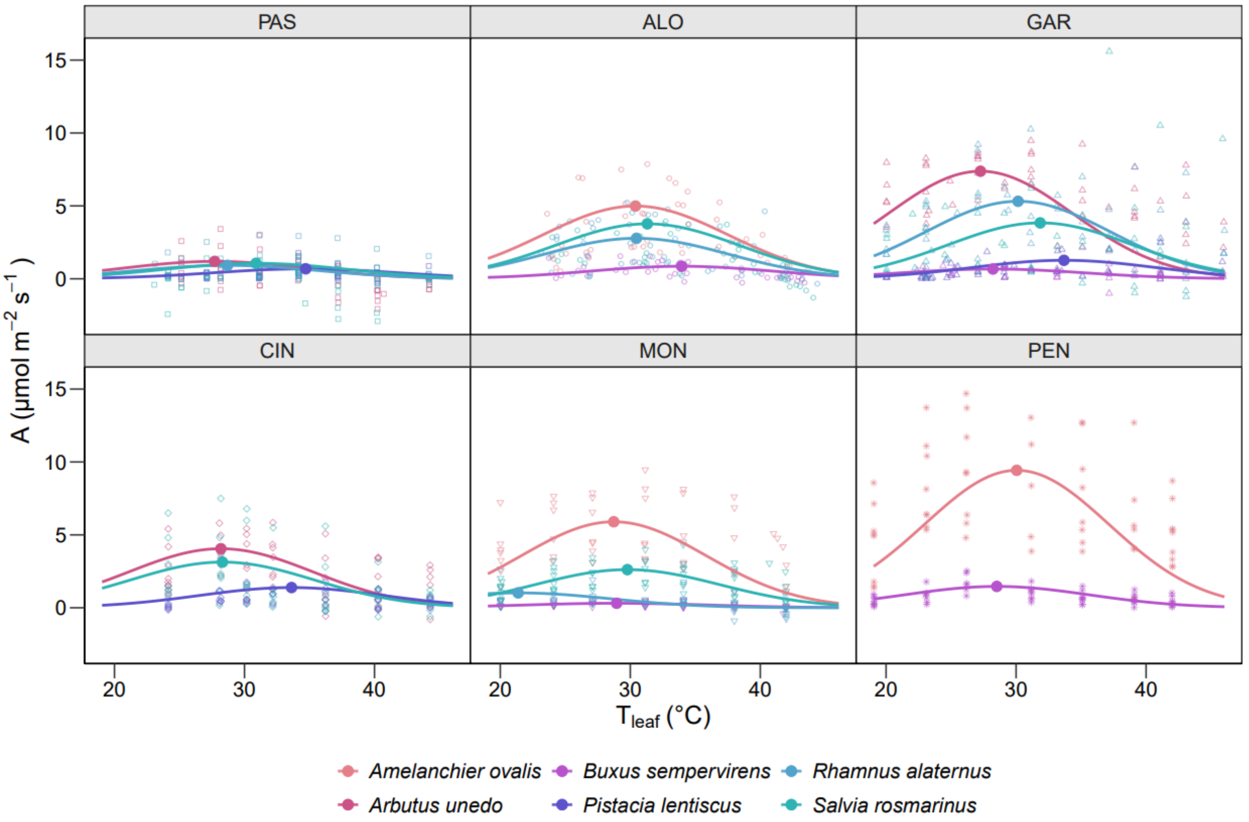


**Figure S4**: Photosynthesis-leaf temperature response curves for each species (i.e., *A. ovalis*, *A. unedo*, *B. sempervirens*, *P. lentiscus*, *R. alaternus*, and *S. rosmarinus*) obtained *in-situ* in each site (i.e., PAS: Pas de l'Ase, ALO: Alòs de Balaguer, GAR: Garraf, CIN: Cingles de Bertí, MON: Montsec, PEN: Pentina). Non-linear regression following a second order Gaussian function for each site and species are shown. Photosynthesis thresholds (i.e., leaf optimal assimilation, $A_{opt}$, for its optimal temperature, $T_{opt}$) for each site and species are indicated by filled symbols.

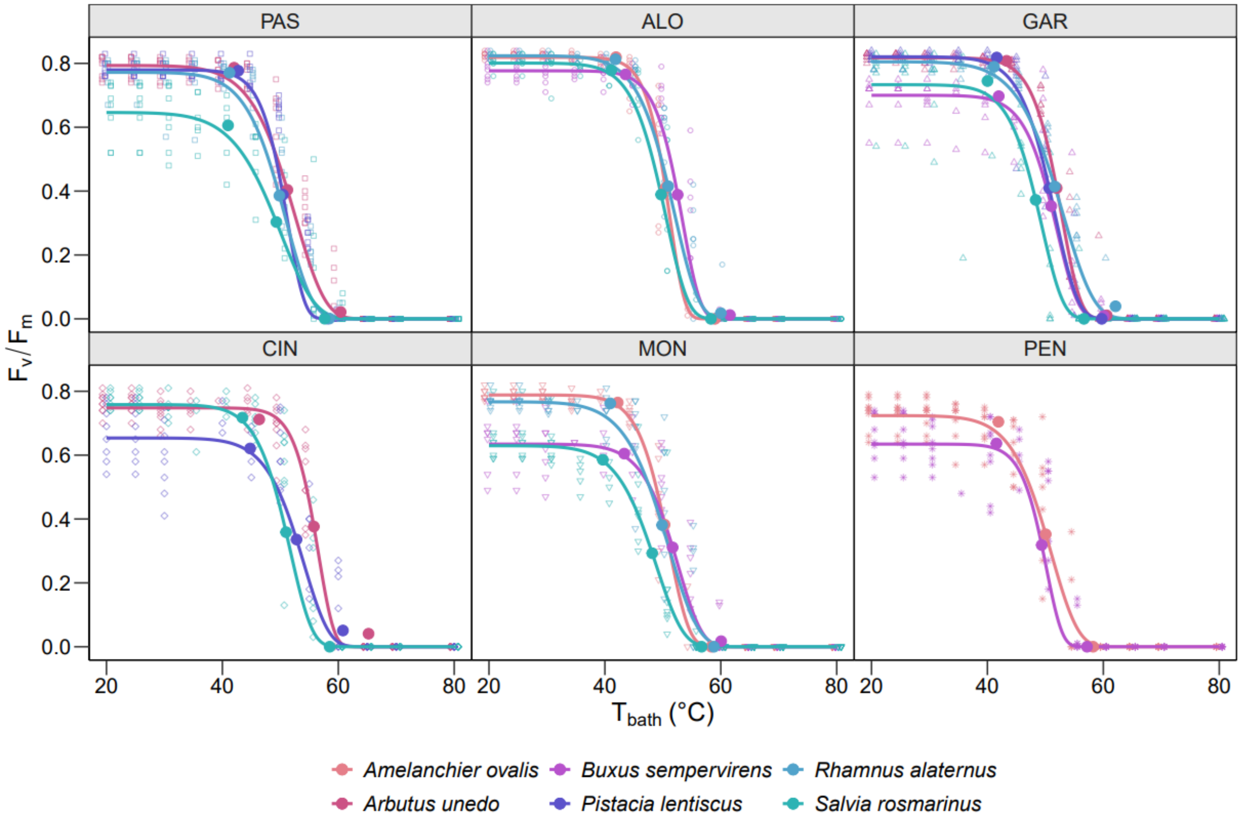


**Figure S5**: Thermotolerance curves showing the photosynthetic efficiency of the photosystem II ($F_v/F_m$) as a function of the water bath temperature for each species (i.e., *A. ovalis*, *A. unedo*, *B. sempervirens*, *P. lentiscus*, *R. alaternus*, and *S. rosmarinus*) from each site (i.e., PAS: Pas de l'Ase, ALO: Alòs de Balaguer, GAR: Garraf, CIN: Cingles de Bertí, MON: Montsec, PEN: Pentina). Sigmoidal Weibull models for each site and species are shown using predicted values. Thermal thresholds (i.e., critical leaf temperature, $T_{crit}$, temperature causing a 50% reduction of $F_v/F_m$, $T_{50}$, maximum temperature, $T_{max}$) for each site and species are indicated by filled symbols.

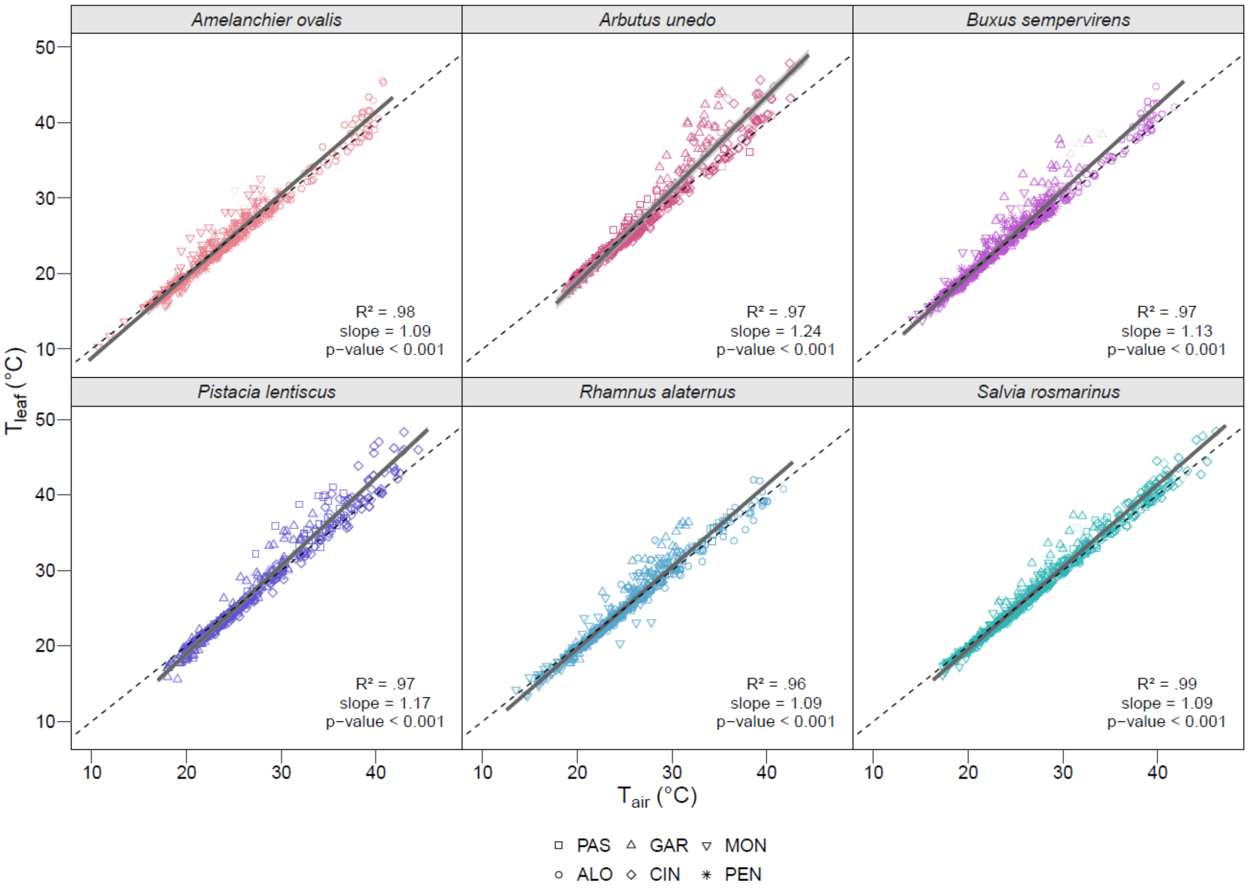


**Figure S6**: Relationship between leaf and air temperature ($T_{leaf}$ and $T_{air}$, respectively) for each species (i.e., *A. ovalis*, *A. unedo*, *B. sempervirens*, *P. lentiscus*, *R. alaternus*, and *S. rosmarinus*) obtained *in-situ* in each site (i.e., PAS: Pas de l'Ase, ALO: Alòs de Balaguer, GAR: Garraf, CIN: Cingles de Bertí, MON: Montsec, PEN: Pentina). The dashed line represents the 1:1 relationship, while the solid grey line represents the linear relationship between $T_{leaf}$ and $T_{air}$ for the corresponding mixed effects models. The $R^2$ indicates the significant relationship, and the slope and p-value indicate the difference to the 1:1 line.

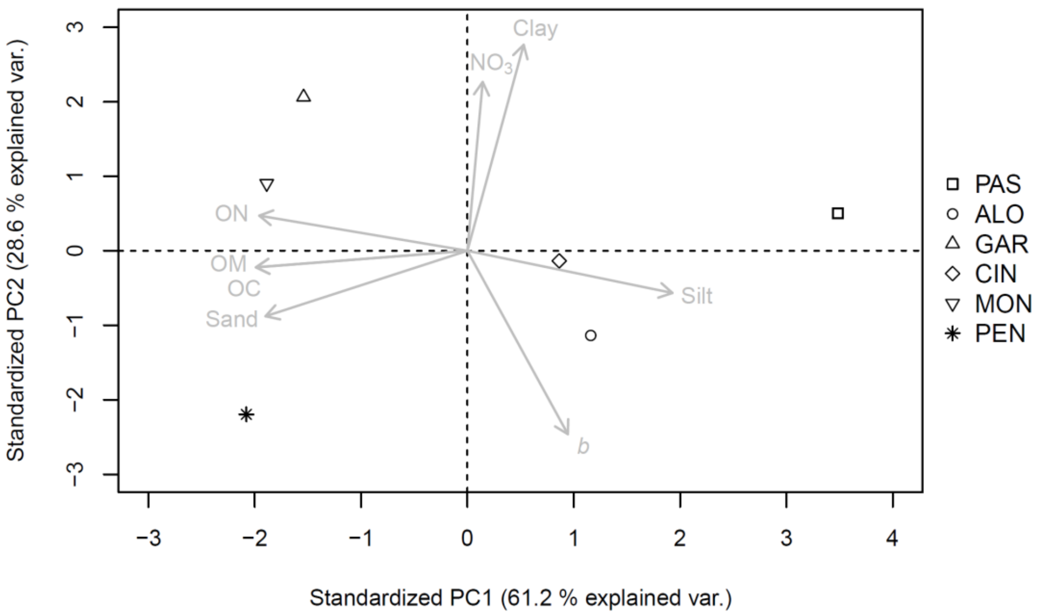


**Figure S7**: Principal components analysis (PCA) of the soil variables, including organic carbon (OC), organic matter (OM), organic nitrogen (ON), nitrate ($NO_3$), clay, silt, sand and *b* coefficient from the Saxton equation (*b*) across all sites (i.e., PAS: Pas de l'Ase, ALO: Alòs de Balaguer, GAR: Garraf, CIN: Cingles de Bertí, MON: Montsec, PEN: Pentina).

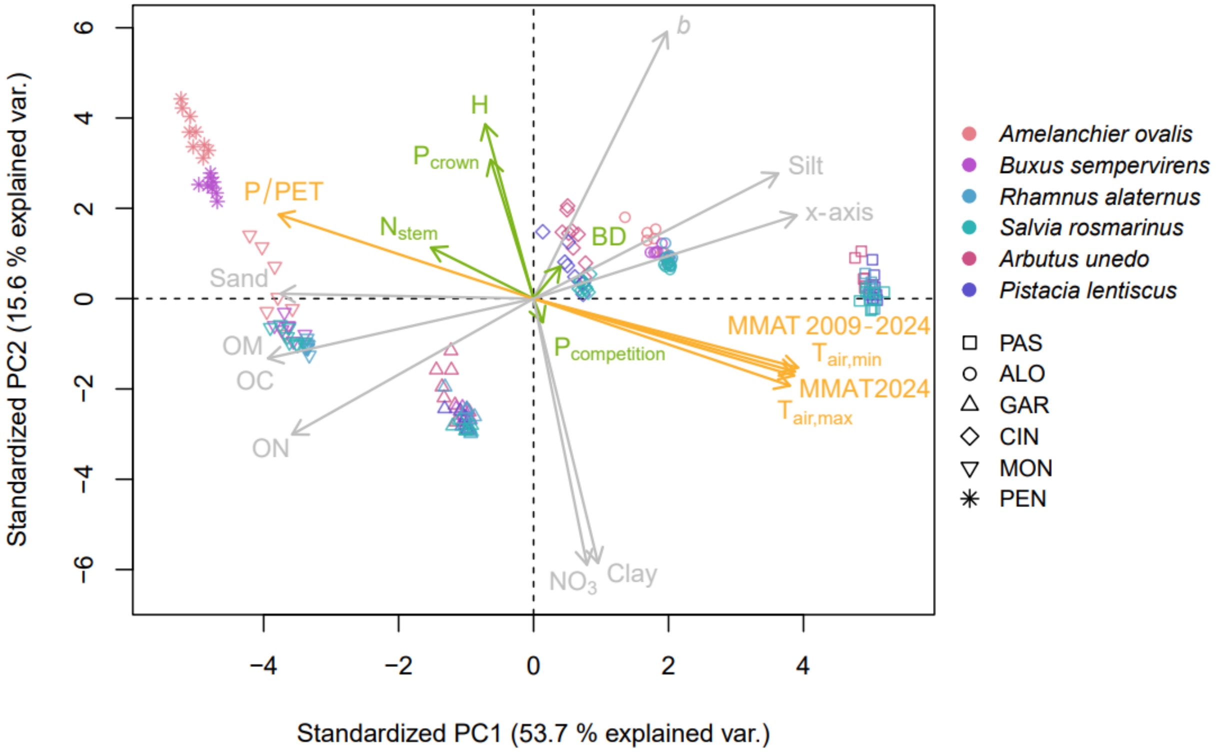


**Figure S8**: Principal components analysis (PCA) of the models fixed effects, including the climatic variables (i.e., maximum monthly air temperature, MMAT 2009-2024; MMAT 2024; air maximum temperature, $T_{air,max}$; air minimum temperature, $T_{air,min}$; P/PET; orange), the soil variables (i.e., *b* coefficient from the Saxton equation, *b*; x-axis from the soil PCA; organic carbon, OC; organic matter, OM; organic nitrogen, ON; nitrate, $NO_3$; clay; silt; sand; grey), the individuals variables (i.e., height, H; basal diameter, BD; stem number, $N_{stem}$; plant competition, $P_{competition}$; plant crown, $P_{crown}$; green) across all species (i.e., *A. ovalis*, *A. unedo*, *B. sempervirens*, *P. lentiscus*, *R. alaternus*, and *S. rosmarinus*), and sites (i.e., PAS: Pas de l'Ase, ALO: Alòs de Balaguer, GAR: Garraf, CIN: Cingles de Bertí, MON: Montsec, PEN: Pentina). Variables were all standardized and log-transformed whenever required to satisfy normality assumptions.

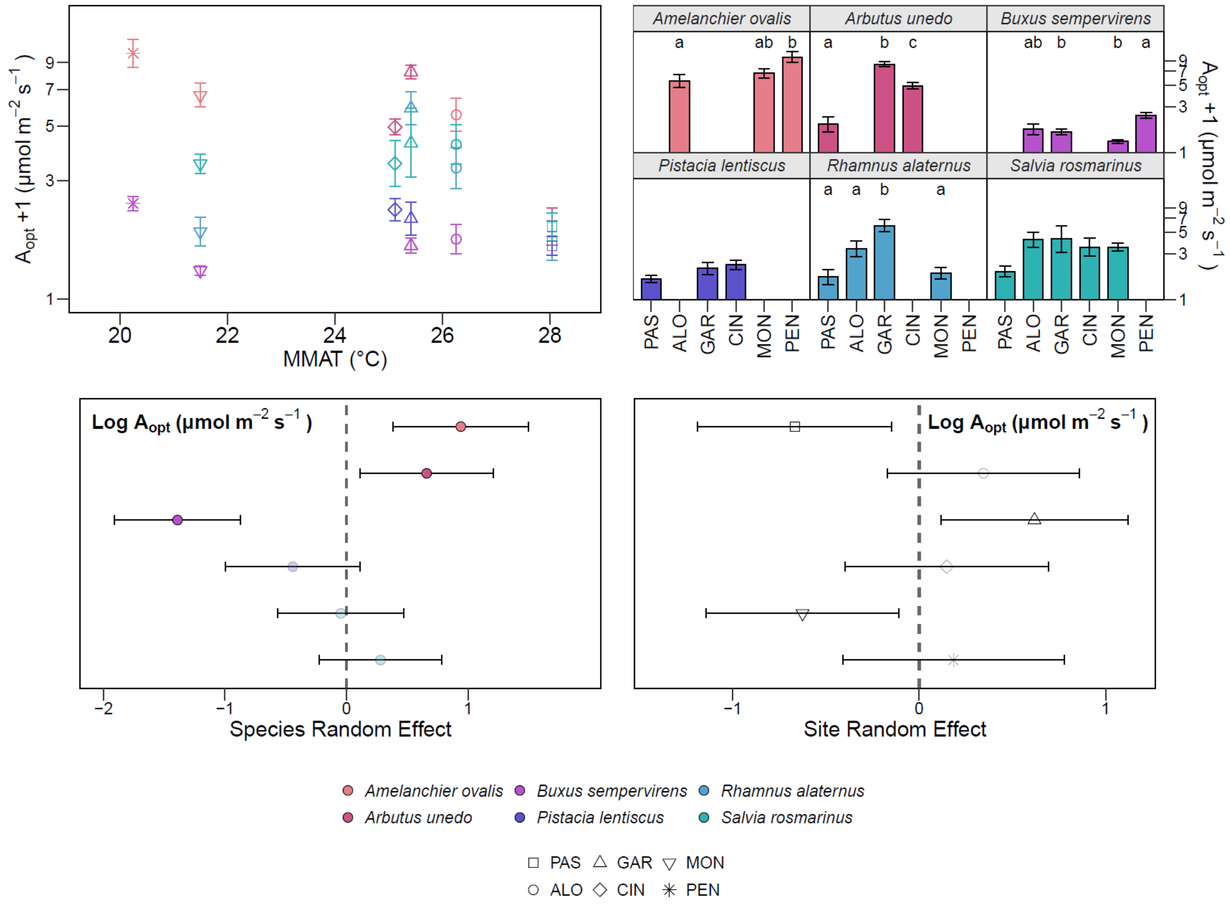


**Figure S9**: (a) Leaf optimal assimilation ($A_{opt}$) as a function of the maximum monthly air temperature (MMAT) for 2024; (b) $A_{opt}$ for each site (i.e., PAS: Pas de l'Ase, ALO: Alòs de Balaguer, GAR: Garraf, CIN: Cingles de Bertí, MON: Montsec, PEN: Pentina). Significant differences between sites are highlighted for each species with letters (Tukey' HSD post hoc test, alpha = 0.05); and (c, d) species and site random effect of the linear mixed-effects models on $A_{opt}$. Significant effects are highlighted for each species and site by symbol' opacity (confidence intervals ≠ 0). Color represents the species (i.e., *A. ovalis*, *A. unedo*, *B. sempervirens*, *P. lentiscus*, *R. alaternus*, and *S. rosmarinus*). Shape represents the sites.

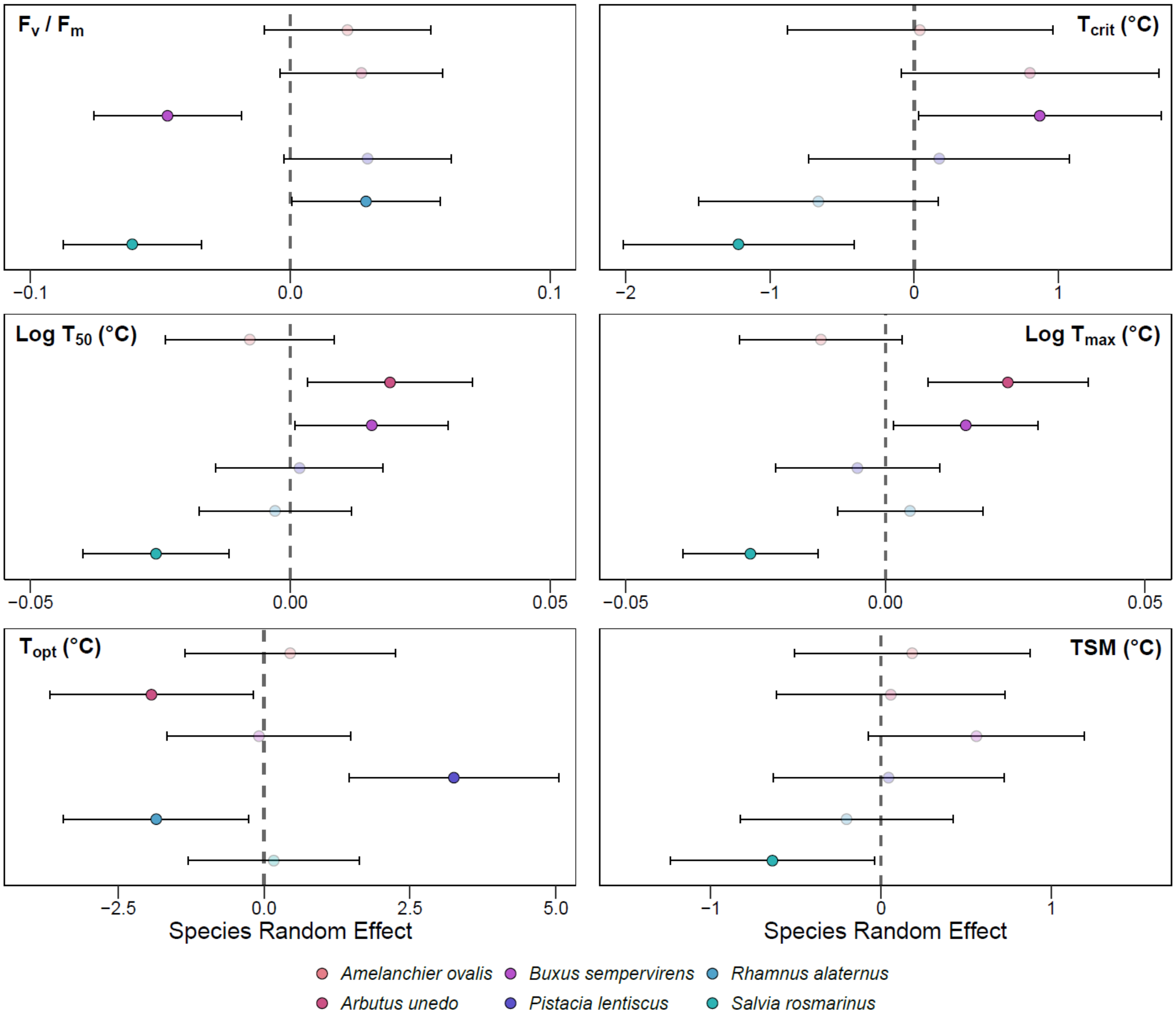


**Figure S10**: Species random effect of the quadratic or linear mixed-effects models on the photosynthetic efficiency of the photosystem II ($F_v/F_m$), critical leaf temperature ($T_{crit}$), temperature causing a 50% reduction of $F_v/F_m$ ($T_{50}$), maximum temperature ($T_{max}$), leaf optimal temperature ($T_{opt}$), and leaf thermal safety margin (TSM) for *A. ovalis* (light pink; *n* = 24 shrubs), *A. unedo* (pink; *n* = 25 shrubs), *B. sempervirens* (magenta; *n* = 32 shrubs), *P. lentiscus* (purple; *n* = 23 shrubs), *R. alaternus* (blue; *n* = 31 shrubs), and *S. rosmarinus* (cyan; *n* = 38; mean ± CI). Significant effects are highlighted for each species by symbol' opacity (confidence intervals ≠ 0).

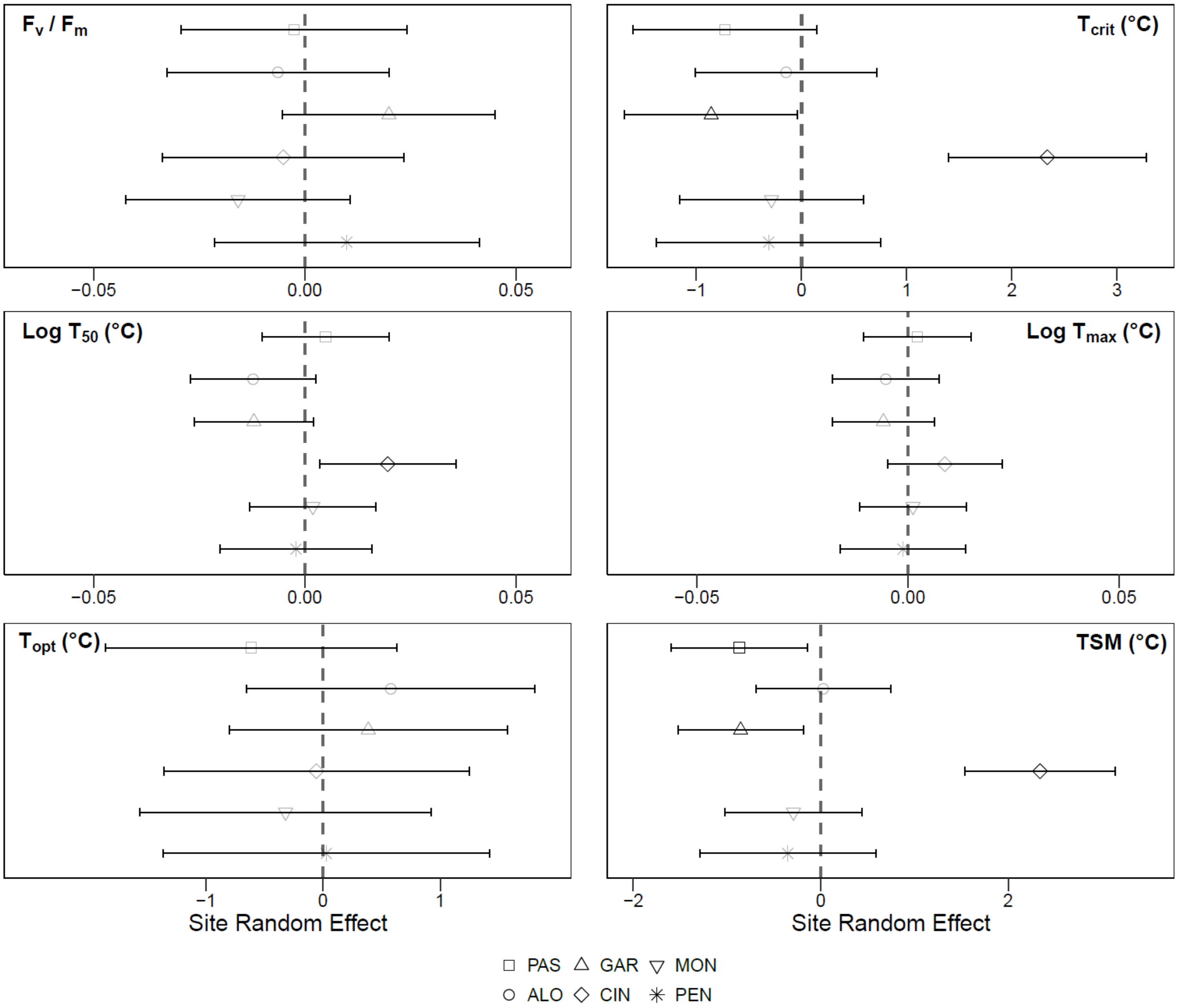


**Figure S11**: Site random effect of the quadratic or linear mixed-effects models on the photosynthetic efficiency of the photosystem II ($F_v/F_m$), critical leaf temperature ($T_{crit}$), temperature causing a 50% reduction of $F_v/F_m$ ($T_{50}$), maximum temperature ($T_{max}$), leaf optimal temperature ($T_{opt}$), and leaf thermal safety margin (TSM) for PAS (Pas de l'Ase, *n* = 31 shrubs), ALO (Alòs de Balaguer, *n* = 32 shrubs), GAR (Garraf, *n* = 38 shrubs), CIN (Cingles de Bertí, *n* = 25 shrubs), MON (Montsec, *n* = 30 shrubs), PEN (Pentina, *n* = 17 shrubs; mean ± CI). Significant effects are highlighted for each species by symbol' opacity (confidence intervals ≠ 0).

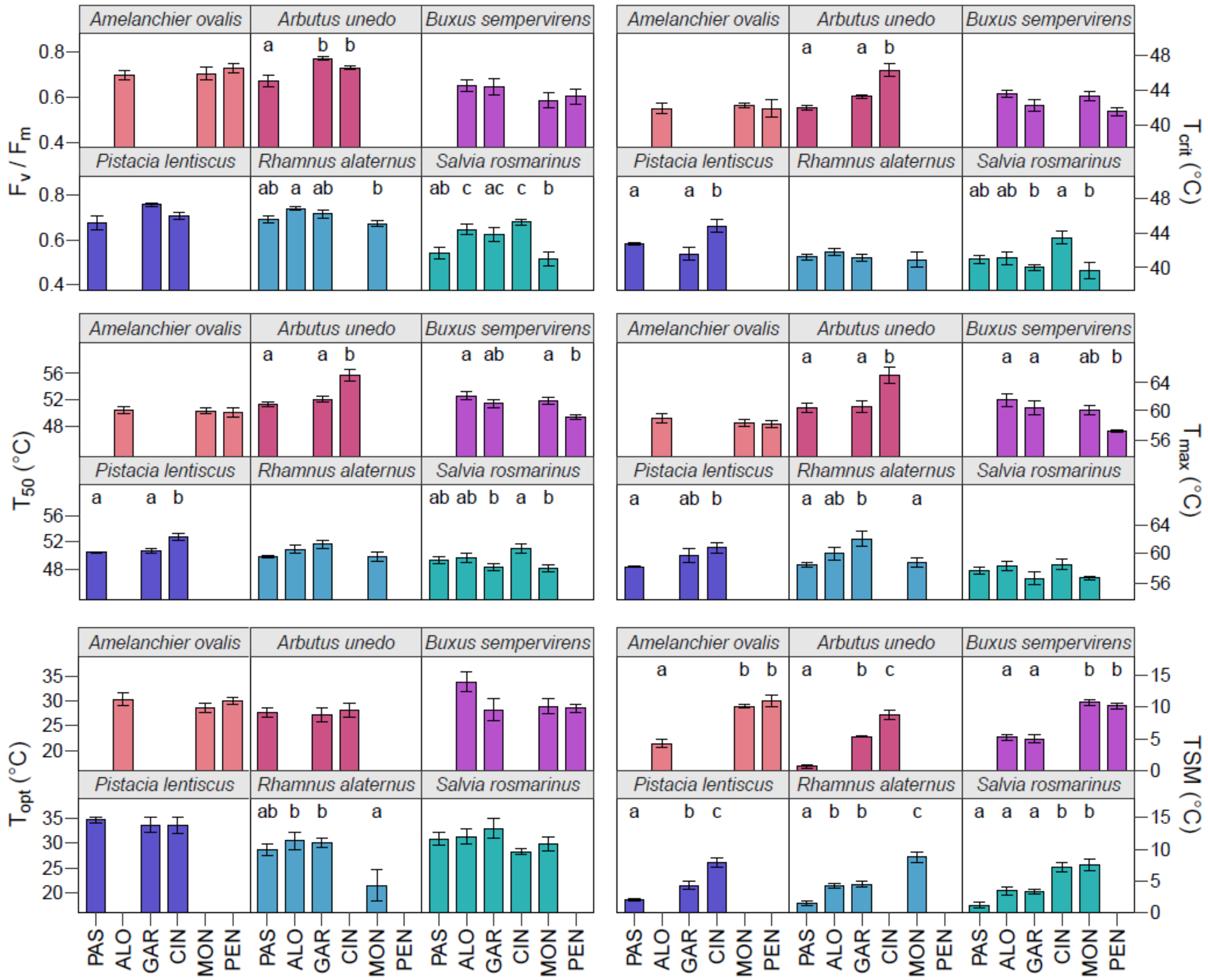


**Figure S12**: Photosynthetic efficiency of the photosystem II ($F_v/F_m$), critical leaf temperature ($T_{crit}$), temperature causing a 50% reduction of $F_v/F_m$ ($T_{50}$), maximum temperature ($T_{max}$), leaf optimal temperature ($T_{opt}$), and leaf thermal safety margin (TSM) for each species (i.e., *A. ovalis*, *A. unedo*, *B. sempervirens*, *P. lentiscus*, *R. alaternus*, and *S. rosmarinus*) and sites (i.e., PAS: Pas de l'Ase, ALO: Alòs de Balaguer, GAR: Garraf, CIN: Cingles de Bertí, MON: Montsec, PEN: Pentina; mean ± CI). Significant differences between sites are highlighted for each species with letters (Tukey' HSD post hoc test, alpha = 0.05).

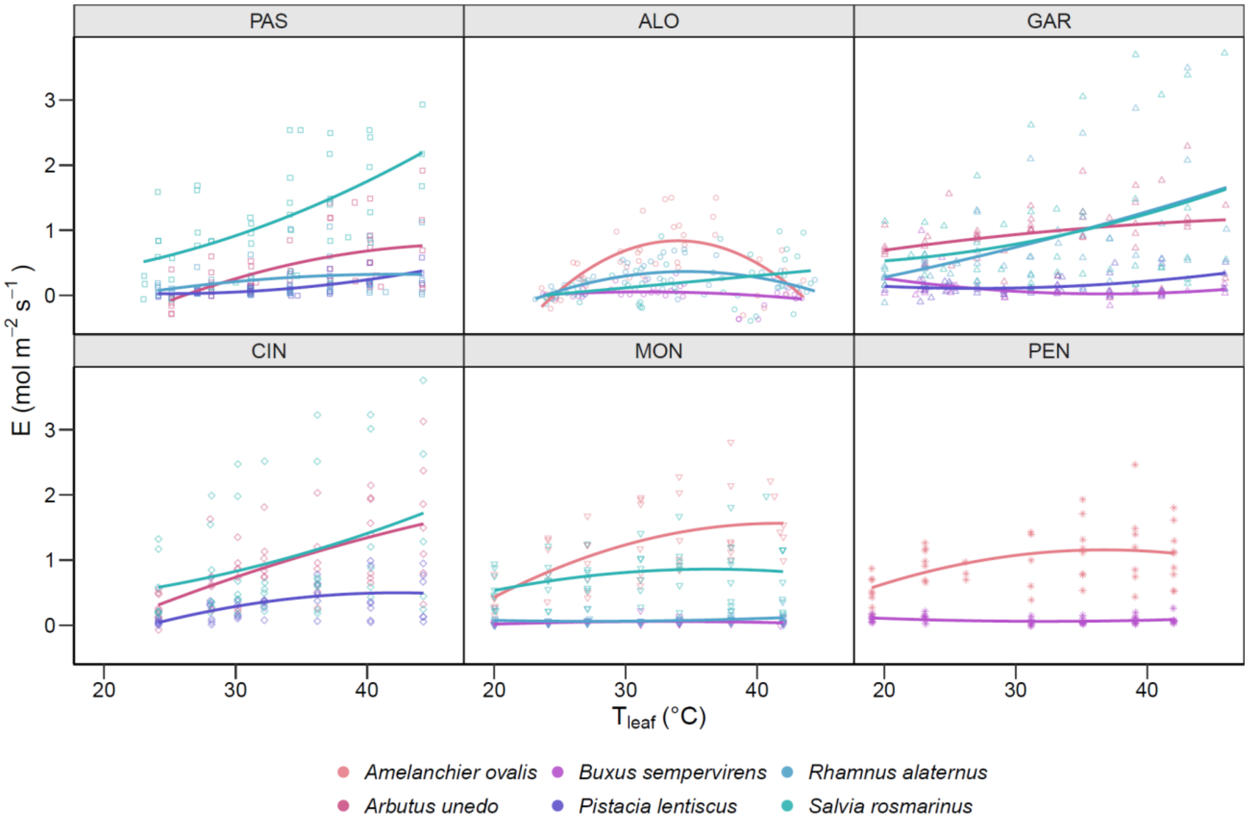


**Figure S13**: Transpiration-leaf temperature response curves for each species (i.e., *A. ovalis*, *A. unedo*, *B. sempervirens*, *P. lentiscus*, *R. alaternus*, and *S. rosmarinus*) obtained *in-situ* in each site (i.e., PAS: Pas de l'Ase, ALO: Alòs de Balaguer, GAR: Garraf, CIN: Cingles de Bertí, MON: Montsec, PEN: Pentina). Non-linear regression following a second order Gaussian function are shown.

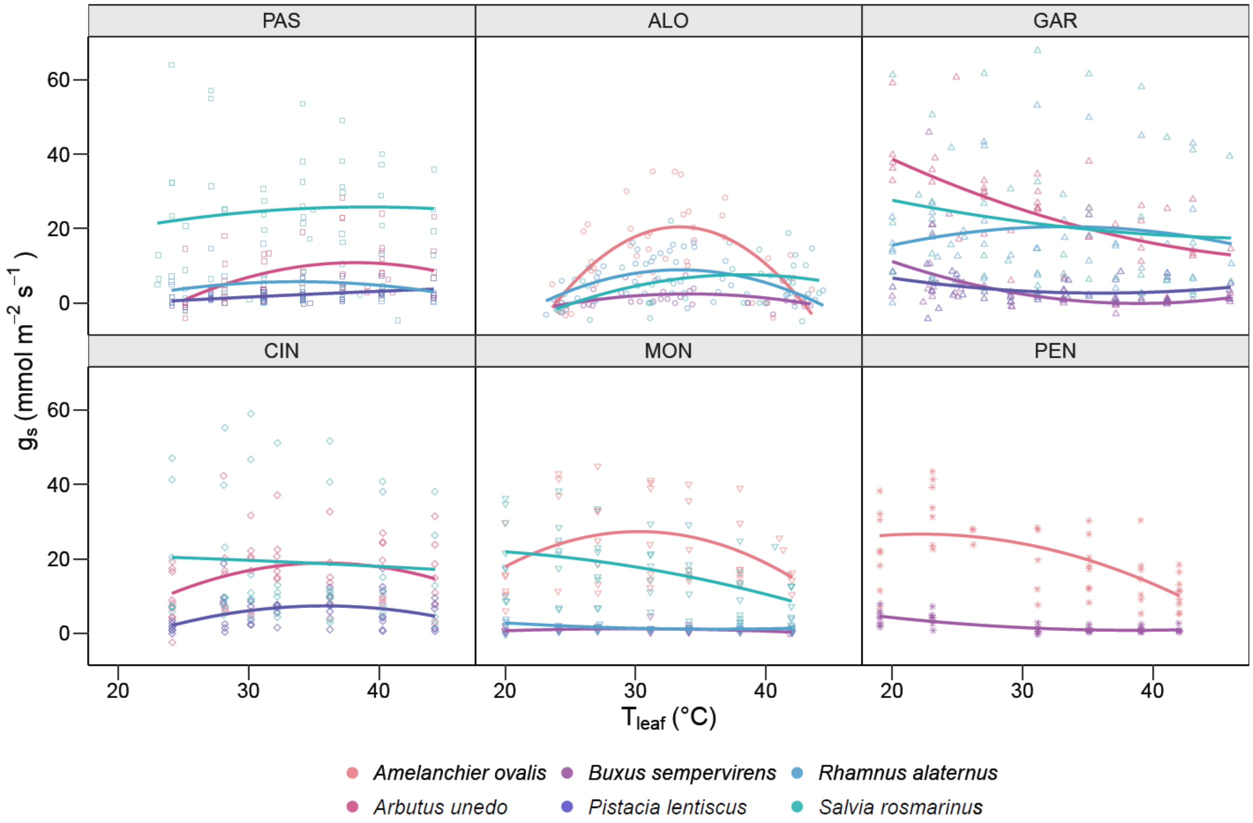


**Figure S14**: Stomatal conductance-leaf temperature response curves for each species (i.e., *A. ovalis*, *A. unedo*, *B. sempervirens*, *P. lentiscus*, *R. alaternus*, and *S. rosmarinus*) obtained *in-situ* in each site (i.e., PAS: Pas de l'Ase, ALO: Alòs de Balaguer, GAR: Garraf, CIN: Cingles de Bertí, MON: Montsec, PEN: Pentina). Non-linear regression following a second order Gaussian function are shown.

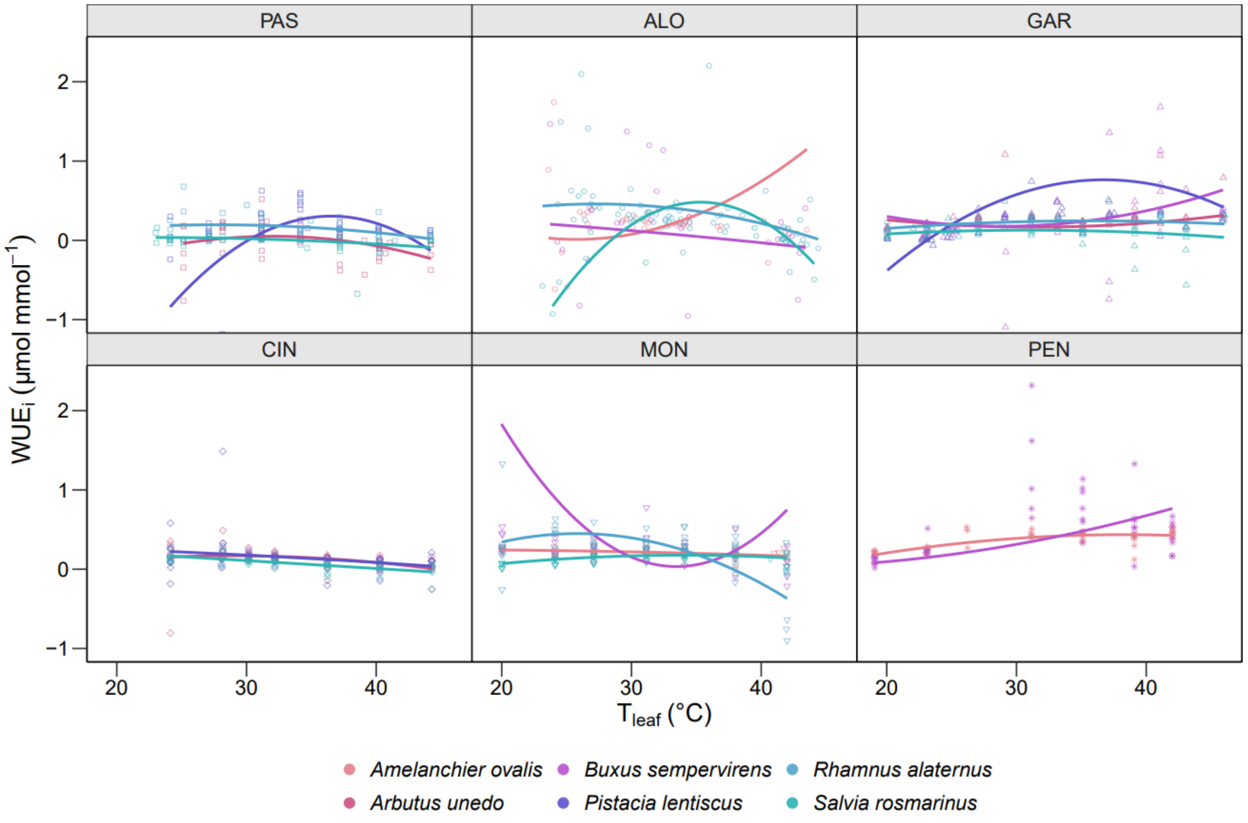


**Figure S15**: Intrinsic water-use efficiency-leaf temperature response curves for each species (i.e., *A. ovalis*, *A. unedo*, *B. sempervirens*, *P. lentiscus*, *R. alaternus*, and *S. rosmarinus*) obtained *in-situ* in each site (i.e., PAS: Pas de l'Ase, ALO: Alòs de Balaguer, GAR: Garraf, CIN: Cingles de Bertí, MON: Montsec, PEN: Pentina). Non-linear regression following a second order Gaussian function are shown.

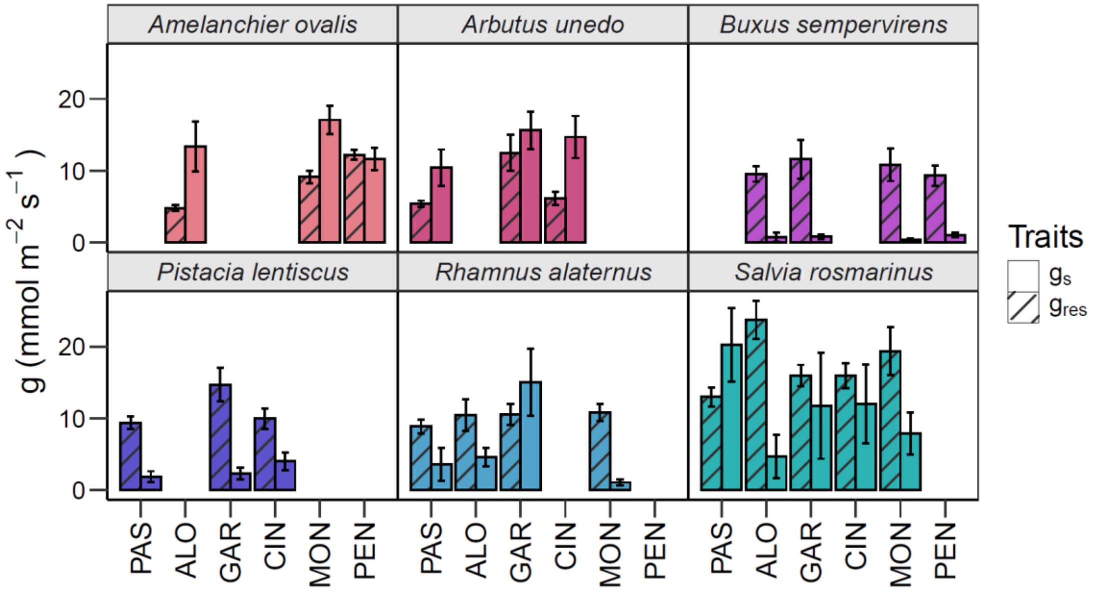


**Figure S16**: Leaf stomatal conductance ($g_s$) measured at the highest leaf temperature and leaf residual conductance ($g_{res}$) measured at 25°C for each species (i.e., *A. ovalis*, *A. unedo*, *B. sempervirens*, *P. lentiscus*, *R. alaternus*, and *S. rosmarinus*) and sites (i.e., PAS: Pas de l'Ase, ALO: Alòs de Balaguer, GAR: Garraf, CIN: Cingles de Bertí, MON: Montsec, PEN: Pentina; mean ± CI).

**Methods S1**: Leaf residual conductance ($g_{res}$) determination.

Leaf residual conductance ($g_{res}$) was measured on one excised sunlight leaf per individuals adjacent to those measured for $F_v/F_m$, leaf gas exchange temperature response curves, $T_{leaf}$ and $T_{air}$, and thermotolerance curves. Leaves were harvested at dawn (between 05:00 and 06:00 local time), when the stomata were assumed to still be closed. They were placed in a sealed plastic bag, and stored in the dark until to be back to the laboratory. Leaves were rehydrated in a dark and cool room (approx. 25°C) for a maximum of 15 h until the next days' analysis. Leaf area was scan using a flatbed scanner (Epson Perfection V39 II, Amsterdam, Netherlands). The mass-loss curves were obtained by sealing the petiole with melted candle wax and attaching it to a laboratory stand with tape, ensuring a stable air temperature (approx. 25 °C). Every 10 min, leaves were weighted using a high-precision scale. This procedure was repeated at least seven times, and $g_{res}$ was obtained from the slope of the linear relationship between leaf mass and drying time (Pearcy *et al.*, 2000).
